# Biosynthesis of β-(1→5)-Galactofuranosyl Chains of Fungal-Type and O-Mannose-Type Galactomannans within the Invasive Pathogen *Aspergillus fumigatus*

**DOI:** 10.1101/814756

**Authors:** Yuria Chihara, Yutaka Tanaka, Minoru Izumi, Daisuke Hagiwara, Akira Watanabe, Kaoru Takegawa, Katsuhiko Kamei, Nobuyuki Shibata, Kazuyoshi Ohta, Takuji Oka

**Affiliations:** Department of Applied Microbial Technology, Faculty of Biotechnology and Life Science, Sojo University, Kumamoto, Japan; Department of Infection and Host Defense, Tohoku Medical and Pharmaceutical University, Sendai, Japan; Graduate School of Environmental and Life Science, Okayama University, Okayama, Japan; Medical Mycology Research Center, Chiba University, Chiba, Japan; Department of Bioscience and Biotechnology, Faculty of Agriculture, Kyushu University, Fukuoka, Japan

**Keywords:** *Aspergillus fumigatus*, cell wall, glycosyltransferase, galactomannan, galactofuranose, glycosylation

## Abstract

The pathogenic fungus *Aspergillus fumigatus* contains galactomannans localized on the surface layer of its cell walls, which are involved in various biological processes. Galactomannans comprise α-(1→2)-/α-(1→6)-mannan and β-(1→5)-/β-(1→6)-galactofuranosyl chains. We previously revealed that GfsA is a β-galactofuranoside β-(1→5)-galactofuranosyltransferase involved in the biosynthesis of β-(1→5)-galactofuranosyl chains. Here, we clarified the entire biosynthesis of β-(1→5)-galactofuranosyl chains in *A. fumigatgus*. Two paralogs exist within *A. fumigatus*: GfsB and GfsC. We show that GfsB and GfsC, in addition to GfsA, are β-galactofuranoside β-(1→5)-galactofuranosyltransferases by biochemical and genetic analyses. GfsA, GfsB, and GfsC can synthesize β-(1→5)-galactofuranosyl oligomers up to lengths of 7, 3, and 5 galactofuranoses within an established *in vitro* highly efficient assay of galactofuranosyltransferase activity. Structural analyses of galactomannans extracted from the strains Δ*gfsB*, Δ*gfsC*, Δ*gfsAC*, and Δ*gfsABC* revealed that GfsA and GfsC synthesized all β-(1→5)-galactofuranosyl residues of fungal-type and O-mannose-type galactomannans, and GfsB exhibited limited function in *A. fumigatus*. The loss of β-(1→5)-galactofuranosyl residues decreased the hyphal growth rate and conidia formation ability as well as increased the abnormal hyphal branching structure and cell surface hydrophobicity, but this loss is dispensable for sensitivity to antifungal agents and virulence toward immune-compromised mice.

**IMPORTANCE:** β-(1→5)-galactofuranosyl residues are widely distributed in the subphylum Pezisomycotina of the phylum Ascomycota. Pezizomycotina includes many plant and animal pathogens. Although the structure of β-(1→5)-galactofuranosyl residues of galactomannans in filamentous fungi was discovered long ago, it remains unclear which enzyme is responsible for biosynthesis of this glycan. Fungal cell wall formation processes are complicated, and information concerning glycosyltransferases is essential for their understanding. In this study, we show that GfsA and GfsC are responsible for the biosynthesis of all β-(1→5)-galactofuranosyl residues of fungal-type and O-mannose-type galactomannans. The data presented here indicates that β-(1→5)-galactofuranosyl residues are involved in cell growth, conidiation, polarity, and cell surface hydrophobicity. Our new understanding of β-(1→5)-galactofuranosyl residue biosynthesis provides important novel insights into the formation of the complex cell wall structure and the virulence of the subphylum Pezisomycotina.

## INTRODUCTION

The cell wall of the pathogenic fungus *Aspergillus fumigatus* comprises several kinds of polysaccharides including chitin, β-(1→3)-glucan, β-(1→3)-/β-(1→4)-glucan, α-(1→3)-glucan, galactosaminogalactan, and galactomannans (GMs) (1–3). These polysaccharides are complexly intertwined to form the three-dimensional structure of cell walls (1, 2). GMs are polysaccharides comprising D-mannose (Man) and D-galactofuranose (Gal*_f_*), localized on the surface layer of cell walls (2), and distinguished into fungal-type galactomannan (FTGM) and O-mannose-type galactomannan (OMGM) (4). FTGM includes core-mannan, a structure wherein α-(1→2)-mannotetraose is linked with α-(1→6)-linkage from 9 to 10, and β-(1→5)-/β-(1→6)-galactofuran side chains (5, 6). FTGM binds to a glycosylphosphatidylinositol anchor for transportation from the Golgi apparatus to the cell surface (7), and the transported FTGM is incorporated into the β-(1→3)-glucan–chitin core of the cell wall by *DFG* family proteins (8). OMGM structure comprises β-(1→5)-/β-(1→6)-galactofuranosyl chains bonded to an O-Man-type glycan with a structure wherein Man is bonded to serine/threonine of a protein as a basic skeleton (6, 9).

Information on GM biosynthesis has been more thoroughly investigated recently (3, 10, 11). CmsA/Ktr4 has been reported to be an α-(1→2)-mannosyltransferase involved in the biosynthesis of the α-(1→2)-mannan backbone of FTGM (12, 13). In the gene-disrupted strain of *cmsA/ktr4* and/or its homolog *cmsB/ktr7*, pronounced hyphal elongation suppression and conidia formation failure were observed (12, 13). Moreover, the Δ*cmsA/ktr4* mutant was significantly less virulent than the parental strain (13). These data indicate that FTGM is crucial for normal cell growth and virulence (12, 13). GfsA is firstly identified as a galactofuranosyltransferase involved in the biosynthesis of OMGM galactofuranosyl residues (14). GfsA is a β-galactofuranoside β-(1→5)-galactofuranosyltransferase also involved in the biosynthesis of FTGM galactofuran side chains (4). However, in the Δ*gfsA* strain of *A. fumigatus*, the β-(1→5)-galactofuranosyl residue was not completely lost (4). The biosynthesis of the remaining β-(1→5)-galactofuranosyl residues remain unclear. Therefore, we focused on clarifying which residual β-(1→5)-galactofuranosyl residues are biosynthesized. There are two paralogs GfsB and GfsC in *A. fumigatus*. We evaluated whether GfsB and GfsC are responsible for biosynthesis of the remaining β-(1→5)-galactofuranosyl residues. We obtained recombinant proteins of GfsA, GfsB, and GfsC to elucidate galactofuranoside chain biosynthesis activity *in vitro* using an established highly efficient assay of galactofuranosyltransferase activity. Furthermore, to investigate the function of *gfs* family proteins *in vivo*, we analyzed the structure of GM extracted from single, double, and triple gene-disruptants of *gfsA*, *gfsB*, and *gfsC*. In this study, we aimed to clarify the biosynthesis and function of β-(1→5)-galactofuranosyl residues in *A. fumigatus*.

## RESULTS

### Features of GfsB and GfsC in A. fumigatus

The Δ*gfsA* disruptant exhibited reduction of the content of β-(1→5)-galactofuranosyl residues within FTGM and OMGM (4); however, these residues remained within the galactomannan fractions. To determine the enzyme synthesizing the remaining β-(1→5)-galactofuranosyl residues, we focused on *gfsA* paralogs, termed *gfsB* (Afu4g13710 in *A. fumigatus*; Af293/AFUB_070620 in *A. fumigatus* A1163) and *gfsC* (Afu4g10170/AFUB_067290). Comparison of cDNA and genome sequences revealed that no introns were present in *gfsB* and *gfsC*, similarly to *gfsA* (4, 14). Analysis of secondary structures using TMHMM revealed that GfsB and GfsC have putative transmembrane domains (GfsB: amino acids 13–35, GfsC: amino acids 23–42) at their *N-*termini, suggesting that both GfsB and GfsC are type II membrane proteins, indicating GfsB and GfsC localization at the Golgi apparatus like GfsA (3, 14). GfsB and GfsC have a conserved metal-binding DXD motif (GfsB: amino acids 237–239, GfsC: amino acids 240–242).

### Enzymatic function of GfsA, GfsB, and GfsC

We previously constructed an *Escherichia coli* strain expressing a recombinant GfsA protein. GfsC was successfully expressed as a soluble protein using a cold-shock expression vector and GfsB was obtained as a soluble fused NusA protein using an *E. coli* expression system. Recombinant 6× His-tagged GfsA, GfsB, and GfsC proteins were purified by Ni^+^ affinity chromatography and analyzed using SDS-PAGE (Fig. S1). The NusA tag of GfsB was cleaved with a HRV 3C protease and removed by Ni-agarose. GfsA, GfsB, and GfsC were visualized as bands close to their predicted respective molecular weights of 57.9, 50.3, and 52.0 kDa. For the galactofuranosyltransferase assay it is essential to use UDP-Gal*_f_* as a sugar donor; however, it is difficult to obtain as it is not commercially available. Thus, we biochemically synthesized UDP-Gal*_f_* using Glf, a UDP-galctopyranose (Gal*_p_*) mutase derived from *E. coli*, followed by HPLC purification (15, 16). This purified UDP-Gal*_f_* was used for the assay within our previous study (4, 14). Because enzymatic equilibrium of the reversible enzyme Glf is inclined to be >93% of the mixture (16), generating much UDP-Gal*_f_* was difficult. To solve this problem, we attempted to improve the galactofuranosyltransferase assay (Fig. 1A). When a small amount of UDP-Gal*_f_* generated is consumed by galactofuranosyltransferase, Glf regenerates UDP-Gal*_f_* to maintain equilibrium (Fig. 1A). Glf oxidizes FADH_2_ to FAD when converting UDP-Gal*_p_* to UDP-Gal*_f_*. Therefore, reducing FAD to FADH_2_ is essential for a continuous reaction (Fig. 1A). Sodium dithionite (SD) was used for re-reduction of FADH_2_ from FAD. Dithionite ion plays a role as a driving horse for proceeding to galactofuranosylation by FADH_2_ reduction from FAD within the continuous reaction, which continues until UDP-Gal*_p_* is almost lost. Based on this principle, we developed a highly efficient assay for galactofuranosyltransferase activity using Glf and Gfs proteins (Fig. 1A). Chemically synthesized 4-methylumbelliferyl-β-D-galactofuranoside (4MU-β-D-Gal*_f_*) or *p*-nitrophenyl-β-D-galactofuranoside (pNP-β-D-Gal*_f_*) was used as acceptor substrate (17, 18). When commercially available pNP-β-D-Gal*_f_* was used as an acceptor substrate instead of 4MU-β-D-Gal*f*, pNP-β-D-Gal*_f_* was not detected by UV300 absorbance, suggesting pNP-β-D-Gal*_f_* variance within the structure by SD (19) (Fig. S2 (d) and (e)). Although NADH/NADPH instead of SD could be used as a reducing reagent against FAD, SD promoted galactofuranosyltransferase reaction more effectively than NADH/NADPH (Fig. S2 (c) and (f)).

**Figure 1.**
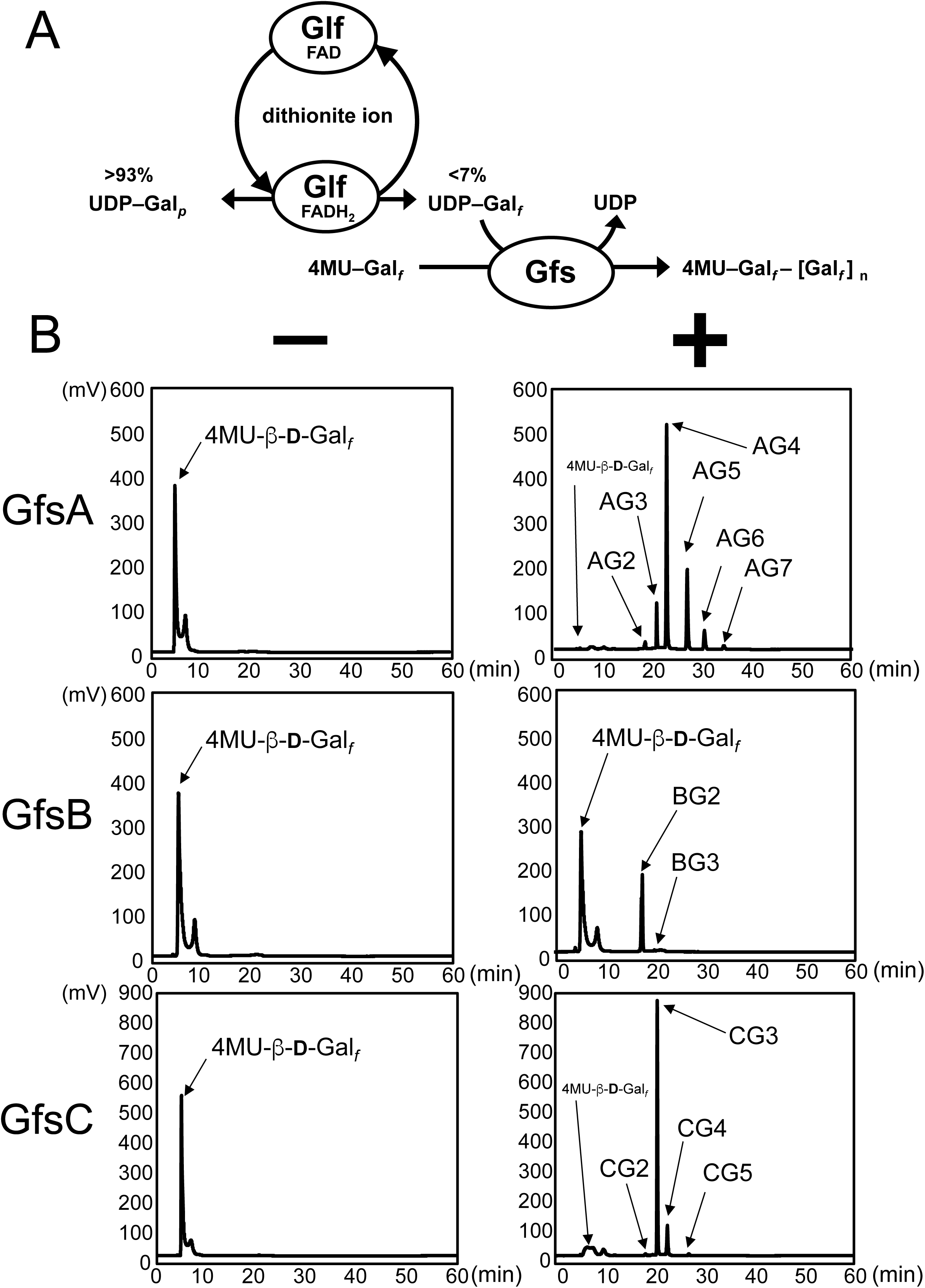
*In vitro* method for measuring galactofuranosyltransferase activity using a continuous reaction. (A) Schematic diagram of continuous reaction in galactofuranosyltransferase activity assay. Glf is UDP-galctopyranose (Gal*_p_*) mutase derived from *E. coli* to generate UDP-Gal*_f_* from UDP-Gal*_p_*. The enzymatic equilibrium of a reversible Glf enzyme is skewed >93% to UDP-Gal*_p_*. When a small amount of UDP-Gal*_f_* generated is consumed by galactofuranosyltransferase, Glf reverts to UDP-Gal*_f_* to maintain equilibrium. Glf oxidizes FADH_2_ to FAD when converting from UDP-Gal*_p_* to UDP-Gal*_f_*. Therefore, reducing FAD to FADH_2_ is essential for a continuous reaction. The dithionite ion plays a role as a driving horse for proceeding to galactofuranosylation by reduction of FADH_2_ from FAD within the continuous reaction. (B) Chromatograms of *in vitro* assay of galactofuranosyltransferase activity of GfsA, GfsB, and GfsC. Enzyme activities were assayed as described in the Materials and Methods. 4.5 µg of purified GfsA, GfsB, or GfsC were used as enzyme. Chromatograms indicate typical results of the assay with/without GfsA, GfsB, or GfsC (*right and left panels, respectively*). The assays lacking GfsA, GfsB, or GfsC showed no novel peak generation (*left panels*), but in contrast, fractions containing GfsA, GfsB, or GfsC did have new products (defined as AG2-AG7 for GfsA; BG2 and BG3 for GfsB; CG2-CG5 for GfsC; *right panels*). Retention times were 18.0 min for AG2, BG2, and CG2, 20.4 min for AG3, BG3, and CG3, 23.4 min for AG4 and CG4, 26.5 min for AG5 and CG5, 30.2 for AG6, and 34.3 min for AG7. Gal*_f_*, galactofuranose; Gal*_p_*, galactopyranose; Glf, UDP-galactopyranose mutase from *E. coli*; UDP, uridine diphosphate; FAD, flavin adenine dinucleotide; 4MU, 4-methylumbelliferyl.

GfsA-lacking fraction showed no new peak generation, but fractions with GfsA showed six new peaks at 18.0, 20.4, 23.4, 26.5, 30.2, and 34.3 min (defined as AG2–AG7, respectively; Fig. 1B, *upper panels*). GfsB-lacking fraction showed no new peak generation, but fractions with GfsB had two new peaks at 18.0 and 20.4 min (BG2 and BG3, respectively; Fig. 1B, *middle panels*). GfsC-lacking fraction showed no new peak generation, but fractions with GfsC had four new peaks at 18.0, 20.4, 23.4, and 26.5 min (CG2–CG5, respectively; Fig. 1B, *bottom panels*). Table 1 shows the mass-to-charge ratios (*m/z*) of enzymatic products of GfsA, GfsB, and GfsC as identified by LC/MS. The differences of each peak were calculated as 162.1, indicating that a hexose molecule is continuously attached and each peak was identical to the theoretical molecular mass of the molecule sequentially added to 4MU-β-D-Gal*_f_* by Gal*_f_*.

**Table 1.**
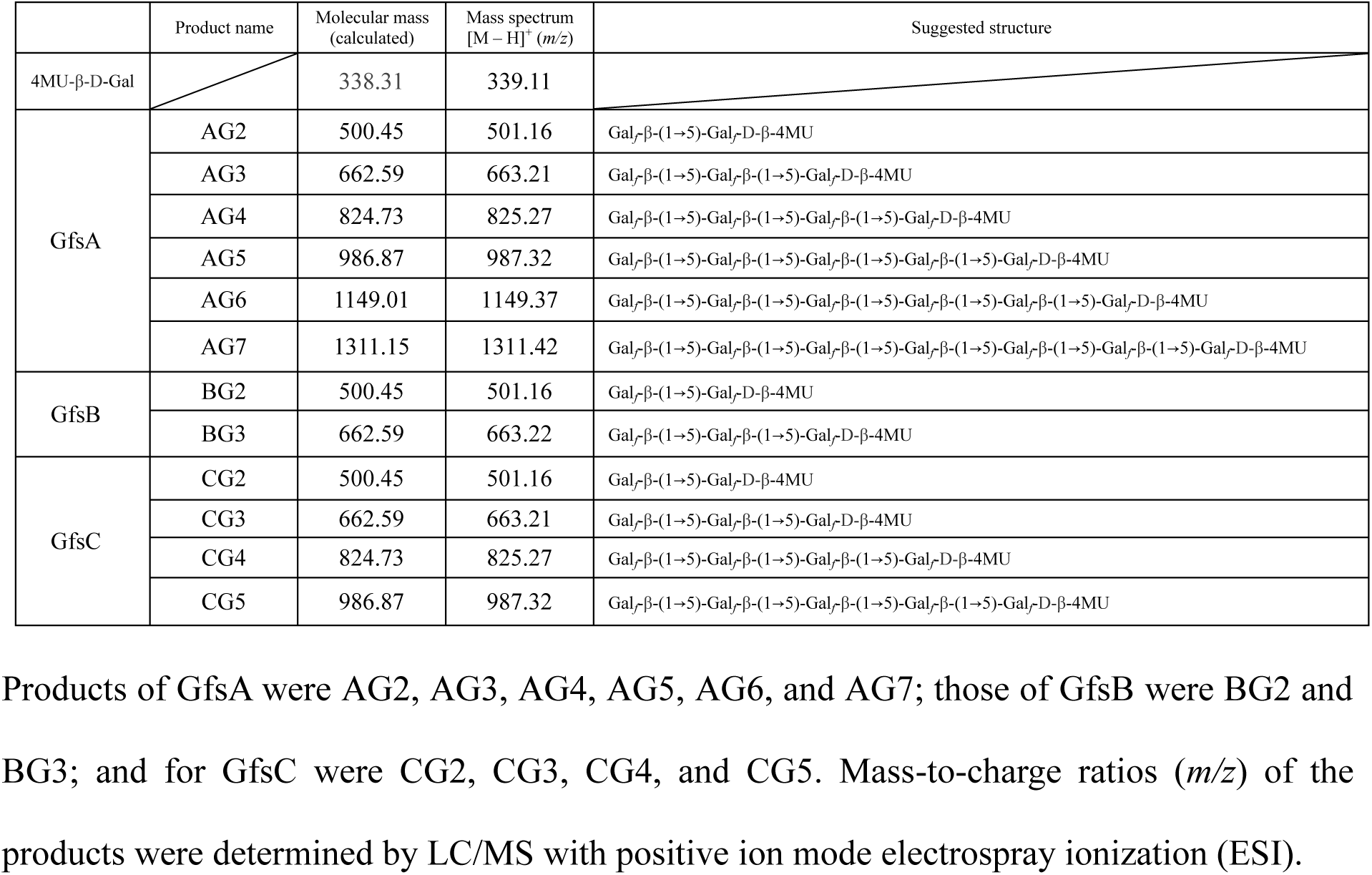
List of mass-to-charge ratios (m/z) of enzymatic products of GfsA, GfsB, and GfsC identified by LC/MS.

To further determine chemical structure, we collected >1 mg of AG3, BG2, and CG3 with HPLC and analyzed the sample using ^1^H-NMR (Fig. 2) with 4MU-β-D-Gal*_f_* as a control. Chemical shift values for H-1 position of the Gal*_f_* residue in t-Gal*_f_*-β-(1→5)-Gal*_f_*-β-(1→5)-Gal*_f_*-D-β-4MU structures are 5.22 (signal A), 5.20 (signal B), and 5.79 (signal C) ppm from the non-reducing end, according to previous reports (4, 6). Signals for AG3, BG2, and CG3 were in agreement with the reported chemical shift values (Fig. 2). To obtain further evidence for glycosidic linkage, we collected 500 μg of AG3, AG4, BG2, CG3, and CG4 using HPLC and analyzed the sample using methylation analysis for each compound. This sample was methylated then hydrolyzed, and subsequently analyzed by GC-MS (Fig. 3). The retention times for t-Gal*_f_*→, 5-Gal*_f_*1→ and 6-Gal*_f_*1→ were 16.36, 18.40, and 19.56 min, respectively, under these analysis conditions (6, 20). AG3, AG4, BG2, CG3, and CG4 displayed a peak at 16.36 min (Fig. 3), indicating presence of terminal Gal*_f_* residues. In addition, AG3, AG4, BG2, CG3, and CG4 had a peak at 18.40 but not 19.56 min (Fig. 3), indicating that the all added Gal*_f_* residue was attached to the C-5 position of the first Gal*_f_* residue. These results indicate that GfsB and GfsC are also β-(1→5)-galactofuranosyltransferases and that GfsA, GfsB, and GfsC could not transfer a Gal*_f_* residue to the C-6 position in contrast to GlfT2, the bacterial β-(1→5)-/β-(1→6)-galactofuranosyltransferase (21, 22).

**Figure 2.**
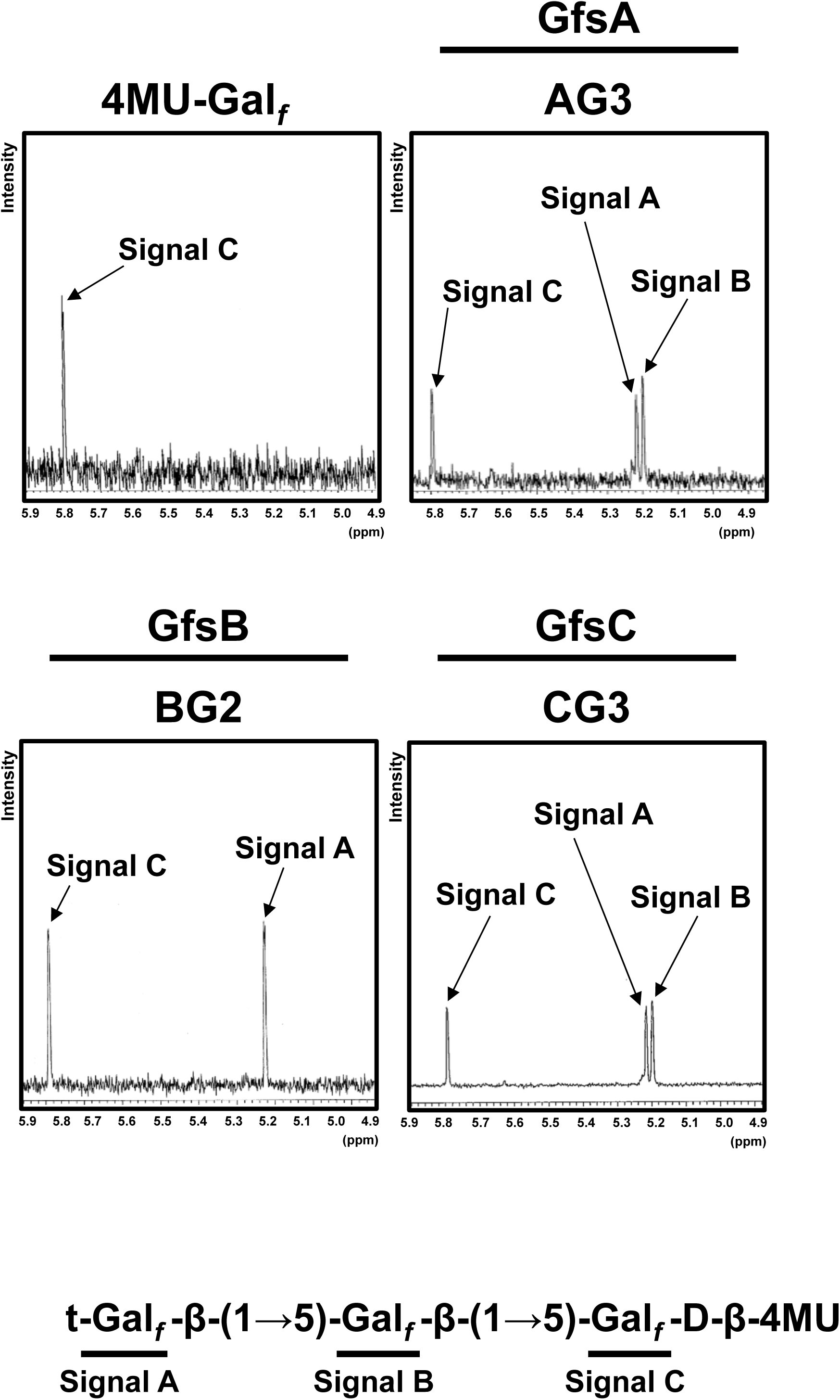
^1^H-NMR analyses of enzymatic products of GfsA, GfsB, and GfsC using 4MU-β-D-Gal*_f_* as an acceptor substrate. ^1^H-NMR charts for 4MU-β-D-Gal*_f_*, AG3, BG2, and CG3. The 5.8 ppm signal was detected in the ^1^H-NMR chart for 4MU-β-D-Gal*_f_*. The chemical shift values of BG2 of the H-1 position of the underlined Gal*_f_* residue in the <u>Gal*_f_*</u>-β-(1→5)-Gal*_f_*-β-4MU and Gal*_f_*-β-(1→5)-<u>Gal*_f_*</u>-β-4MU structures are 5.22 and 5.79 ppm, respectively, according to previous reports (4, 6, 17, 18). The chemical shift values of AG3 and CG3 of the H-1 position of the underlined Gal*_f_* residue in the <u>Gal*_f_*</u>-β-(1→5)-Gal*_f_*-β-(1→5)-Gal*_f_*-β-4MU, Gal*_f_*-β-(1→5)-<u>Gal*_f_*</u>-β-(1→5)-Gal*_f_*-β-4MU and Gal*_f_*-β-(1→5)-Gal*_f_*-β-(1→5)-<u>Gal*_f_*</u>-β-4MU structures are 5.22, 5.20 and 5.79 ppm, respectively, according to previous reports (4, 6, 17, 18).

**Figure 3.**
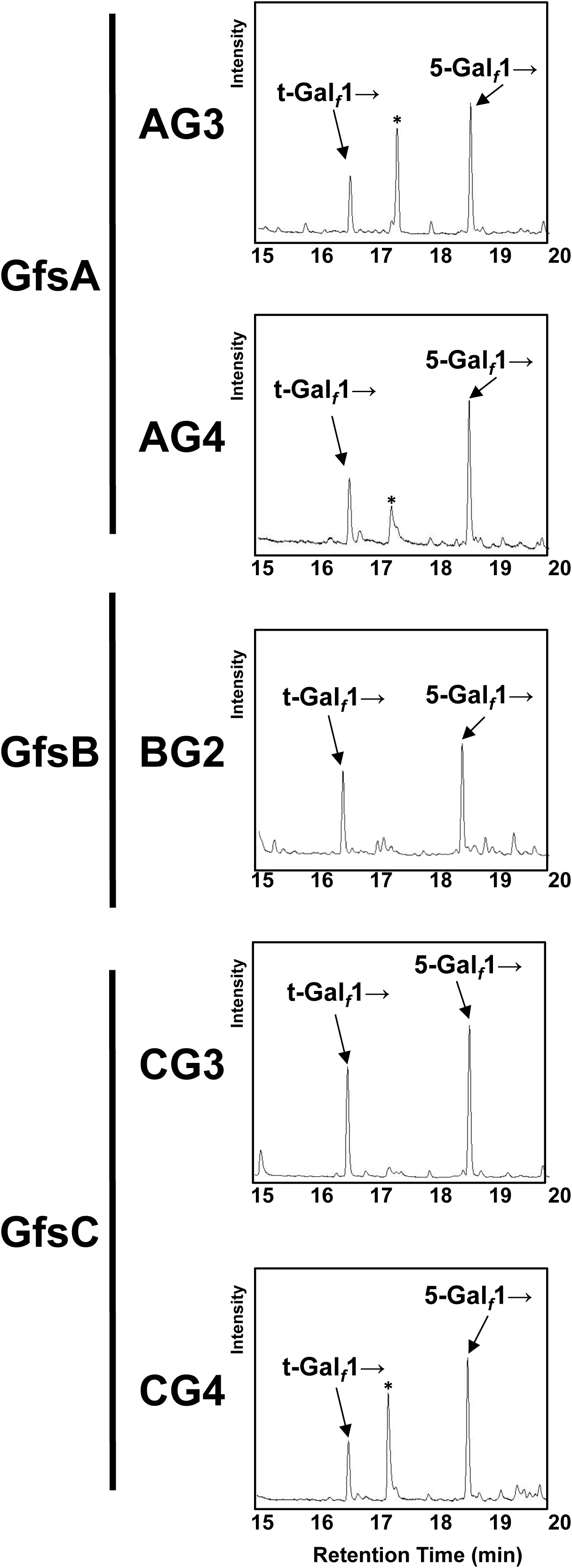
Methylation analyses of enzymatic products of GfsA, GfsB, and GfsC using 4MU-β-D-Gal*f* as an acceptor substrate. 500-µg samples of AG3, AG4, BG2, CG3, and CG4 were analyzed. The retention times under these analysis conditions for t-Gal*_f_*→, Gal*_f_*1→ and 6-Gal*_f_*1→ were 16.36, 18.40, and 19.56 min, respectively (4, 6, 20). The asterisk indicates an artificial peak that appears frequently.

### Role of GfsB and GfsC in GM biosynthesis

To clarify the function of the *gfs* family *in vivo*, we constructed Δ*gfsB*, Δ*gfsC*, Δ*gfsC::C*, Δ*gfsAC*, and Δ*gfsABC* strains (Fig. S3, S4 and S5). To identify the effect of gene disruption on the structure of GMs, those extracted from the mycelia of *A. fumigatus* strains were purified by cetyl trimethyl ammonium bromide precipitation with boric acid buffer. The GMs designated as FTGM+OMGM contain both FTGM and OMGM (4); these were analyzed by ^13^C-NMR spectroscopy (Fig. 4). Signals at 107.87 ppm and 108.70 ppm of the ^13^C-NMR spectra represent the C-1 positions of the underlined Gal*_f_* residue within the structure of -<u>Gal*_f_*</u>-β-(1→5)-Gal*_f_*-(1→ (β-(1→5)-Gal*_f_*) and -<u>Gal*_f_*</u>-β-(1→6)-Gal*_f_*-(1→ (β-1→6)-Gal*_f_*), respectively, as according to previous reports (6, 23). The signal intensity of β-(1→5)-Gal*_f_* was higher than that of β-(1→6)-Gal*_f_* in the ^13^C-NMR chart of A1151-FTGM+OMGM. Within Δ*gfsB*-FTGM+OMGM there was little difference from A1151-FTGM+OMGM (Fig. 4). In contrast, the intensity of signal of β-(1→5)-Gal*_f_* was inversed in the ^13^C-NMR chart of Δ*gfsC*-FTGM+OMGM, indicating that the amount of β-(1→5)-Gal*_f_* was decreased in the FTGM+OMGM fraction of Δ*gfsC* strains (Fig. 4). The signal intensity of β-(1→5)-Gal*_f_* was recovered in the ^13^C-NMR chart of Δ*gfsC::C*-FTGM+OMGM (Fig. 4). Interestingly, signals of β-(1→5)-Gal*_f_* in the ^13^C-NMR chart of Δ*gfsAC-/*Δ*gfsABC*-FTGM+OMGM were not detected, indicating that β-(1→5)-Gal*_f_* disappeared within the FTGM+OMGM fractions of Δ*gfsAC* and Δ*gfsABC* strains. GC-MS analyses of O-methylalditol acetates derived from methylation analyses of FTGM+OMGMs were performed for the A1151, Δ*gfsB*, Δ*gfsC*, Δ*gfsAC*, Δ*gfsABC*, and Δ*gfsC::C* strains (Table 2). The ratio of the 5-O-substituted Gal*_f_* residue (5-Gal*_f_*1→) of Δ*gfsC* (2.16% ± 0.19%) was lower than that of A1151 (16.31% ± 0.84%); however, the ratio of 5-Gal*_f_*1→ of Δ*gfsB* (15.37% ± 0.71%) was comparable with that of A1151 (Table 2). Interestingly, signals for the 5-Gal*_f_*1→ of Δ*gfsAC* or Δ*gfsABC* were not detected within these FTGM+OMGM fractions (Table 2). These results clearly indicate that β-(1→5)-galactofuranosyl residues disappeared within both Δ*gfsAC* and Δ*gfsABC*. Next, the FTGM galactofuran side chain was prepared and separated by gel filtration chromatography to analyze its length (Fig. 5). FTGM+OMGM fractions were treated with 0.15 M of trifluoroacetic acid at 100°C for 15 min. The resultant samples were applied to gel filtration chromatography to separate the obtained galactofuran side chain (Fig. 5). Consequently, sugar chains consisting of up to 6 monosaccharides were detected in the fractions of A1151, Δ*gfsA* and Δ*gfsC* strains (Fig. 5). Conversely, only monosaccharide was detected in the Δ*gfsAC* strain fraction, indicating that elongation by β-(1→5)-galactofuranosyl residues had not occurred (Fig. 5). These observations clearly indicate that all β-(1→5)-galactofuranosyl residues of FTGM and OMGM in *A. fumigatus* are biosynthesized by GfsA and GfsC.

**Figure 4.**
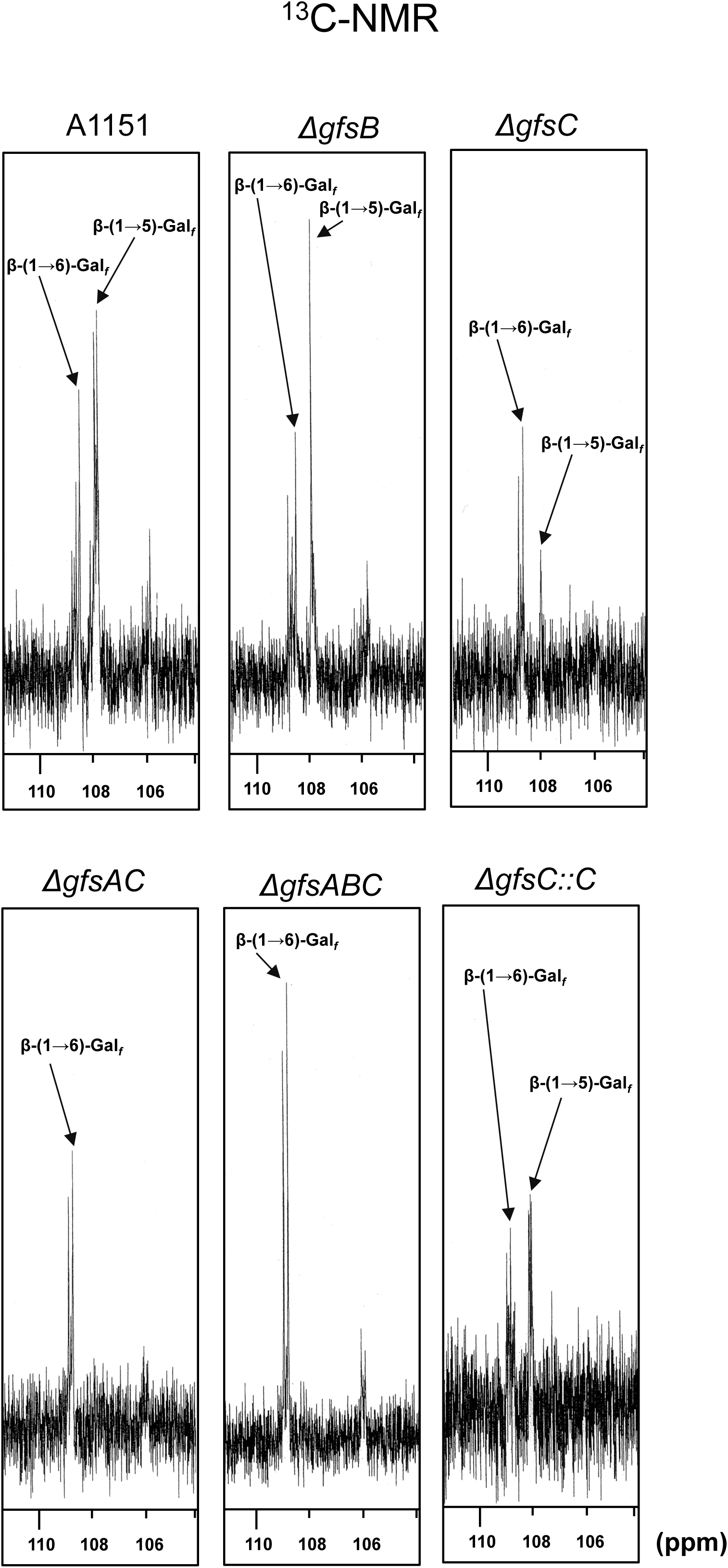
^13^C-NMR analyses of purified FTGM+OMGM fractions from A1151, Δ*gfsB*, Δ*gfsC*, Δ*gfsAB*, Δ*gfsABC*, and Δ*gfsC::C* strains. The signals at 107.87 ppm and 108.70 ppm represent a C-1 chemical shift of the underlined Gal*_f_* residue in the Gal*_f_*-β-(1→5)-<u>Gal*_f_*</u>-β-(1→5)-Gal*_f_*(β-(1→5)-Gal*_f_*) and Gal*_f_*-β-(1→5)-<u>Gal*_f_*</u>-β-(1→6)-<u>Gal*_f_*</u> (β-(1→6)-Gal*_f_*) structures, respectively (6, 20). The carbon chemical shifts were referenced relative to internal acetone at 31.07 ppm, respectively. OMGM, O-mannose-type galactomannan; FTGM, fungal-type galactomannan; FTGM+OMGM, total GM (FTGM + OMGM).

**Figure 5.**
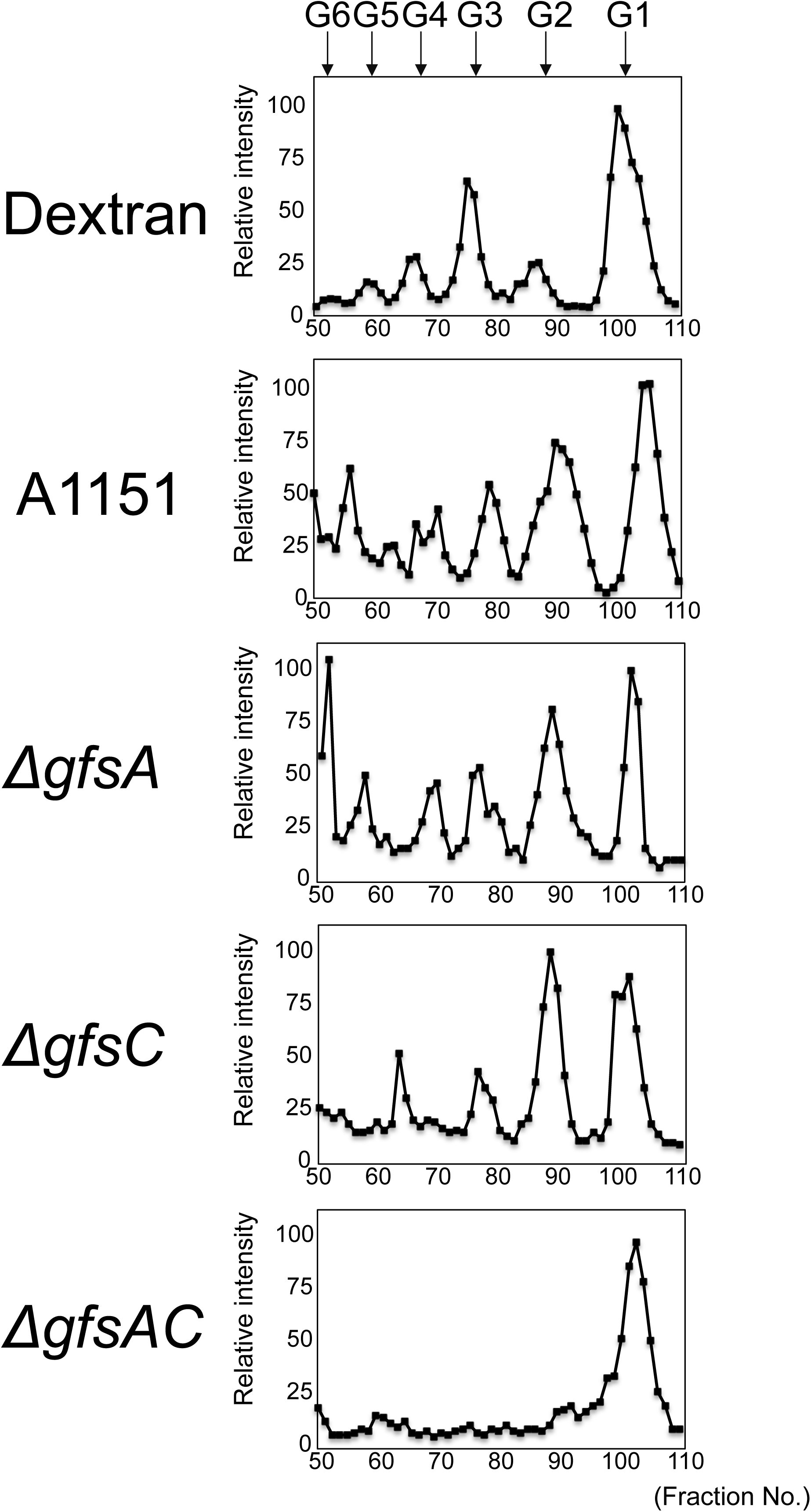
Analysis of galactofuran side chain length of fungal-type galactomannan. Galactofuran side chain was prepared and separated by gel filtration chromatography. FTGM+OMGM fractions were treated with 0.15 M of trifluoroacetic acid at 100°C for 15 min. The resultant samples were applied to gel filtration chromatography using a Bio-Gel P-2 (2 × 90 cm) column and dH_2_O as eluent. The partial acid hydrolysis product of dextran was used as a molecular weight marker. The eluted sugar was detected using the phenol-sulfuric acid method. G1, Glucose; G2, Maltose; G3, Maltotriose; G4, Maltotetraose; G5, Maltopentaose; G6, Maltohexaose; G7, Maltoheptaose.

**Table 2.**
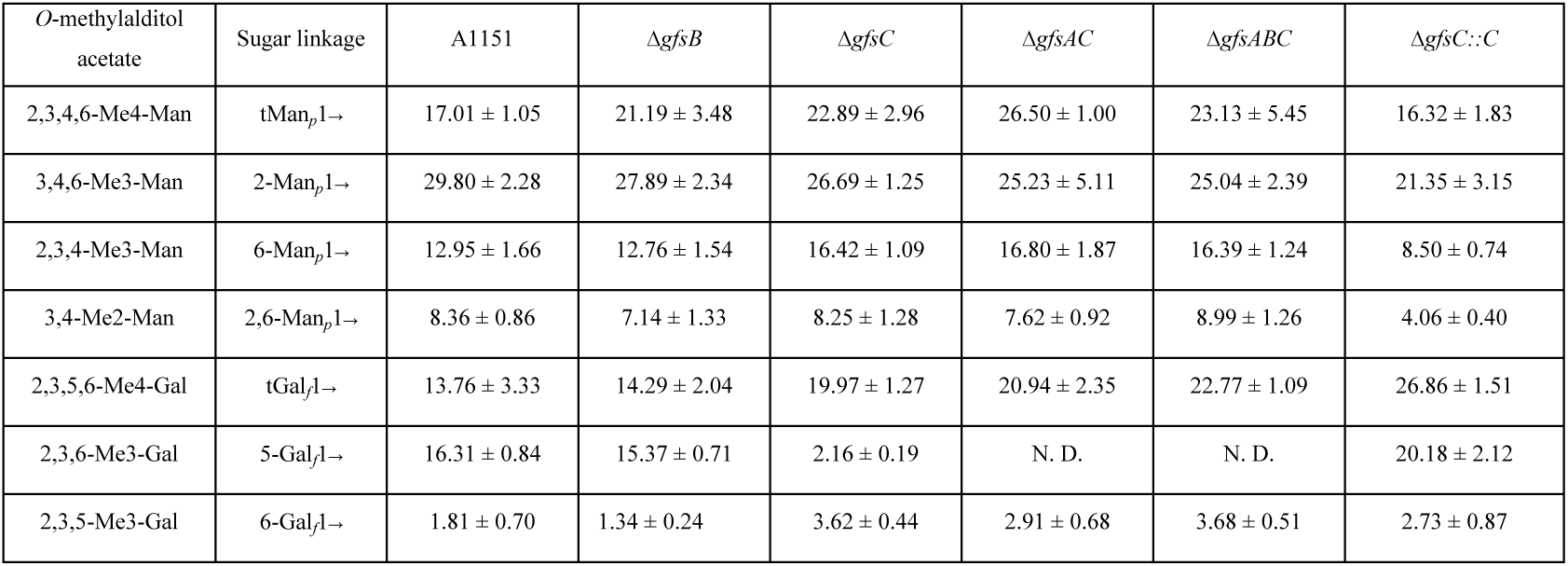
GC-MS analysis of O-methylalditol acetates derived from methylation analyses of galactomannans.

### Phenotypic analyses of disruptant *gfs* family genes

The colony phenotypes of disruptant strains were observed following 3 days of growth at 37°C/50°C on minimal medium (Fig. 6). The colony growth rates of the disruptant strains are shown in Table 3. The colony growth rate of the Δ*gfsA* strain decreased to 85.2% as compared with that of the A1151 strain at 37°C (Table 3). In contrast, colony growth rate percentages of Δ*gfsB* and Δ*gfsC* strains were comparable with that of the A1151 strain at 37°C (Table 3). Growth rate percentages of Δ*gfsAC* and Δ*gfsABC* strains were reduced to 68.4% and 67.8% at 37°C, and to 86.4% and 84.0% at 50°C, respectively (Table 3). When quantifying the number of formed conidia at 37°C, the percentage of the Δ*gfsA* strain decreased to 50.9% compared with that of the A1151 strain. The conidiation efficiencies of Δ*gfsAC* and Δ*gfsABC* were reduced to approximately 32.1% and 25.4% of that of the A1151 strain (Table 4). In contrast, the conidiation efficiency of the Δ*gfsB* and Δ*gfsC* strains did not obviously decrease (Table 4). These results are consistent with the facts that GfsA and GfsC have redundant enzymatic functions and GfsB can only synthesize short β-(1→5)-galactofuranosyl oligomers. These results indicate that the β-(1→5)-galactofuranosyl residues plays an important role in conidia formation and hyphal growth.

**Figure 6.**
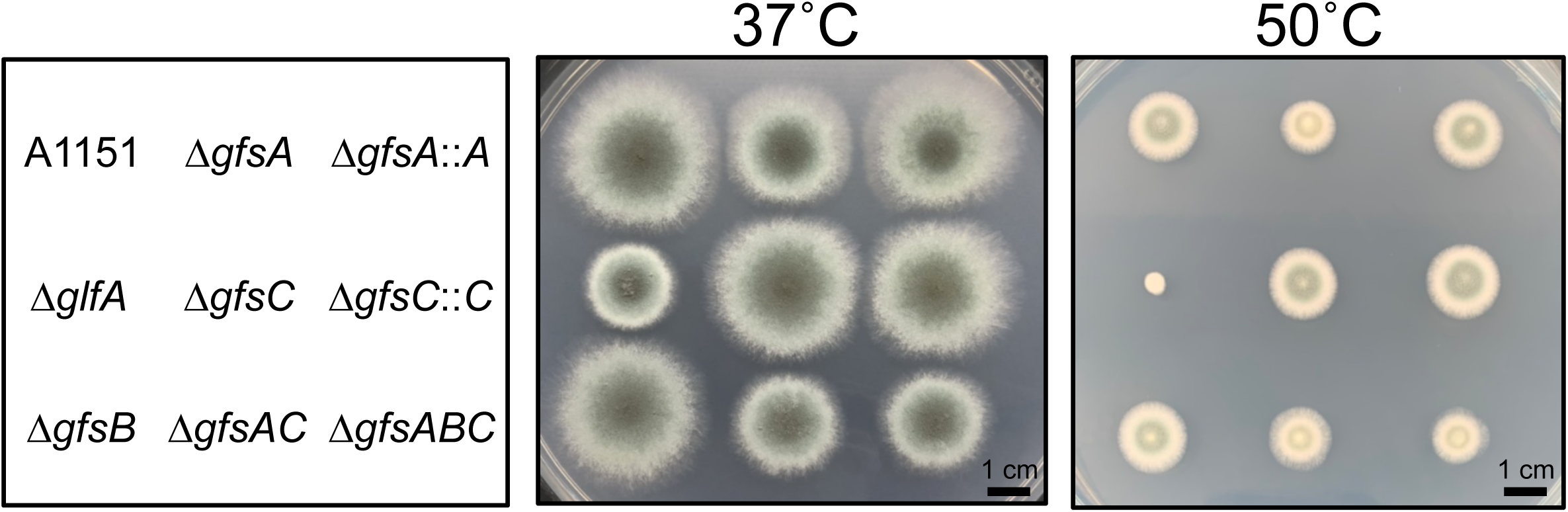
Colony phenotype comparison of A1151, Δ*gfsA*, Δ*gfsB*, Δ*gfsC*, Δ*gfsAB*, Δ*gfsABC*, *gfsA::A* and Δ*gfsC::C* strains. Strain colony images are shown for A1151, Δ*gfsA*, Δ*gfsB*, Δ*gfsC*, Δ*gfsAB*, Δ*gfsABC*, *gfsA::A*, and Δ*gfsC::C*. Conidia were incubated on minimal medium at 37°C (*left*) or 50°C (*right*) for 3 days.

**Table 3.** Colony growth rate of the WT, Δ*gfsA*, Δ*gfsB*, Δ*gfsC*, Δ*gfsAC*, Δ*gfsABC*, Δ*gfsA::A*, andΔ*gfsC::C* strains.

| | A1151 | $\Delta gfsA$ | $\Delta gfsB$ | $\Delta gfsC$ | $\Delta gfsAC$ | $\Delta gfsABC$ | $\Delta gfsA::A$ | $\Delta gfsC::C$ |
| --- | --- | --- | --- | --- | --- | --- | --- | --- |
| 37°C | 0.75 ± 0.06<br>(100%) | 0.63 ± 0.04<br>(85.2%) | 0.73 ± 0.10<br>(98.6%) | 0.76 ± 0.08<br>(102.0%) | 0.51 ± 0.03<br>(68.4%) | 0.50 ± 0.05<br>(67.8%) | 0.78 ± 0.04<br>(104.1%) | 0.76 ± 0.06<br>(101.9%) |
| 50°C | 0.30 ± 0.04<br>(100%) | 0.24 ± 0.03<br>(81.2%) | 0.30 ± 0.03<br>(100.5%) | 0.27 ± 0.11<br>(90.9%) | 0.26 ± 0.03<br>(86.4%) | 0.25 ± 0.03<br>(84.0%) | 0.31 ± 0.03<br>(103.2%) | 0.37 ± 0.03<br>(124.3%) |

**Table 4.** Number of formed conidia of the WT, ΔgfsA, ΔgfsB, ΔgfsC, ΔgfsAC, ΔgfsABC,ΔgfsA::A, andΔgfsC::C strains.

| Strain name | Number of formed conidia (conidia/mm<sup>2</sup>) | Percentage of formed conidia to WT strain |
| --- | --- | --- |
| A1151 | $3.1 \times 10^5 \pm 9.6 \times 10^4$ | 100% |
| $\Delta AfgfsA$ | $1.6 \times 10^5 \pm 9.1 \times 10^3$ | 50.9% |
| $\Delta AfgfsB$ | $3.0 \times 10^5 \pm 5.2 \times 10^4$ | 95.9% |
| $\Delta AfgfsC$ | $2.8 \times 10^5 \pm 3.0 \times 10^4$ | 90.9% |
| $\Delta AfgfsAC$ | $1.0 \times 10^5 \pm 5.7 \times 10^3$ | 32.1% |
| $\Delta AfgfsABC$ | $7.9 \times 10^4 \pm 3.5 \times 10^3$ | 25.4% |
| $\Delta AfgfsA::A$ | $2.6 \times 10^5 \pm 3.5 \times 10^4$ | 82.6% |
| $\Delta AfgfsC::C$ | $3.8 \times 10^5 \pm 3.4 \times 10^4$ | 120.3% |

It was reported that hyphae branching was increasing within the Δ*glfA* strain (24). Therefore, we observed hyphae in Δ*gfsAC* and Δ*gfsABC* strains to determine whether abnormal hyphae branching was also formed in these strains (Fig. 7A). We did observe abnormal hyphae branching at a high frequency, indicating that deficient of β-(1→5)-galactofuranosyl residues causes an increase of abnormal hyphae branching. Reportedly, lack of the Gal*_f_*-containing sugar chains from the cell causes increased cell surface hydrophobicity (24). To confirm increasing cell surface hydrophobicity in Δ*gfsAC* and Δ*gfsABC* strains, whether the amount of adherence of latex beads to the mycelium increased as observed within the Δ*glfA* strain was determined; this adherence was clearly increased (Fig. 7B), indicating that β-(1→5)-galactofuranosyl residues are involved in cell surface hydrophobicity in *A. fumigatus*.

### Sensitivity to antifungal agents and virulence of β-(1→5)-galactofuranosyl residue-deficient strains

Next, we tested the sensitivity of the A1151, Δ*gfsC*, Δ*gfsAC*, and Δ*gfsABC* strains to the widely used clinical antifungal agents micafungin (MCFG), caspofungin (CPFG), amphotericin B (AMPH-B), flucytosine (5-FC), fluconazole (FLCZ), itraconazole (ITCZ), voriconazole (VRCZ), and miconazole (MCZ) (Table 5). Sensitivities of the mutants to antifungal agents were almost identical to those of A1151. The Δ*gfsABC* strain exhibited only slightly greater sensitivity to AMPH-B and MCZ as compared with the A1151 strain (Table 5). We also examined the role of β-(1→5)-galactofuranosyl residues in pathogenesis using a murine infection model (Fig. 8). First, the virulence of A1151, Δ*gfsC* and Δ*gfsC::C* strains were tested within immune-compromised mice. Survival rates did not differ between A1151, Δ*gfsC* and Δ*gfsC::C* infections (Fig. 8A). Virulence of A1151, Δ*gfsC*, Δ*gfsAC*, and Δ*gfsABC* strains were also tested (Fig. 8B). In the aspergillosis model, virulence of the Δ*gfsAC* and Δ*gfsABC* strains were comparable with that of the A1151 strain (Fig. 8B), indicating that a lack of β-(1→5)-galactofuranosyl residues did not influence survival rates of immunosuppressed mice.

**Table 5.** Sensitivity of the WT, Δ*cmsA*, Δ*cmsB*, and Δ*cmsAB* strains to antifungal agents (μg/mL).

|  | MCFG | CPFG | AMPH-B | 5-FC | FLCZ | ITCZ | VRCZ | MCZ |
| --- | --- | --- | --- | --- | --- | --- | --- | --- |
| A1151 | 0.015 | 0.25 | 1 | >64 | >64 | 0.5 | 0.5 | 2 |
| $\Delta AfgfsC$ | 0.015 | 0.25 | 1 | >64 | >64 | 0.5 | 0.5 | 1-2 |
| $\Delta AfgfsAC$ | 0.015 | 0.25 | 1 | >64 | >64 | 0.25-0.5 | 0.5-1 | 1-2 |
| $\Delta AfgfsABC$ | 0.015 | 0.25 | 0.5 | >64 | >64 | 0.5 | 0.5 | 1 |

micafungin (MCFG), caspofungin (CPFG), amphotericin B (AMPH-B), flucytosine (5-FC), fluconazole (FLCZ), itraconazole (ITCZ), voriconazole (VRCZ), and miconazole (MCZ).

**Figure 7.**
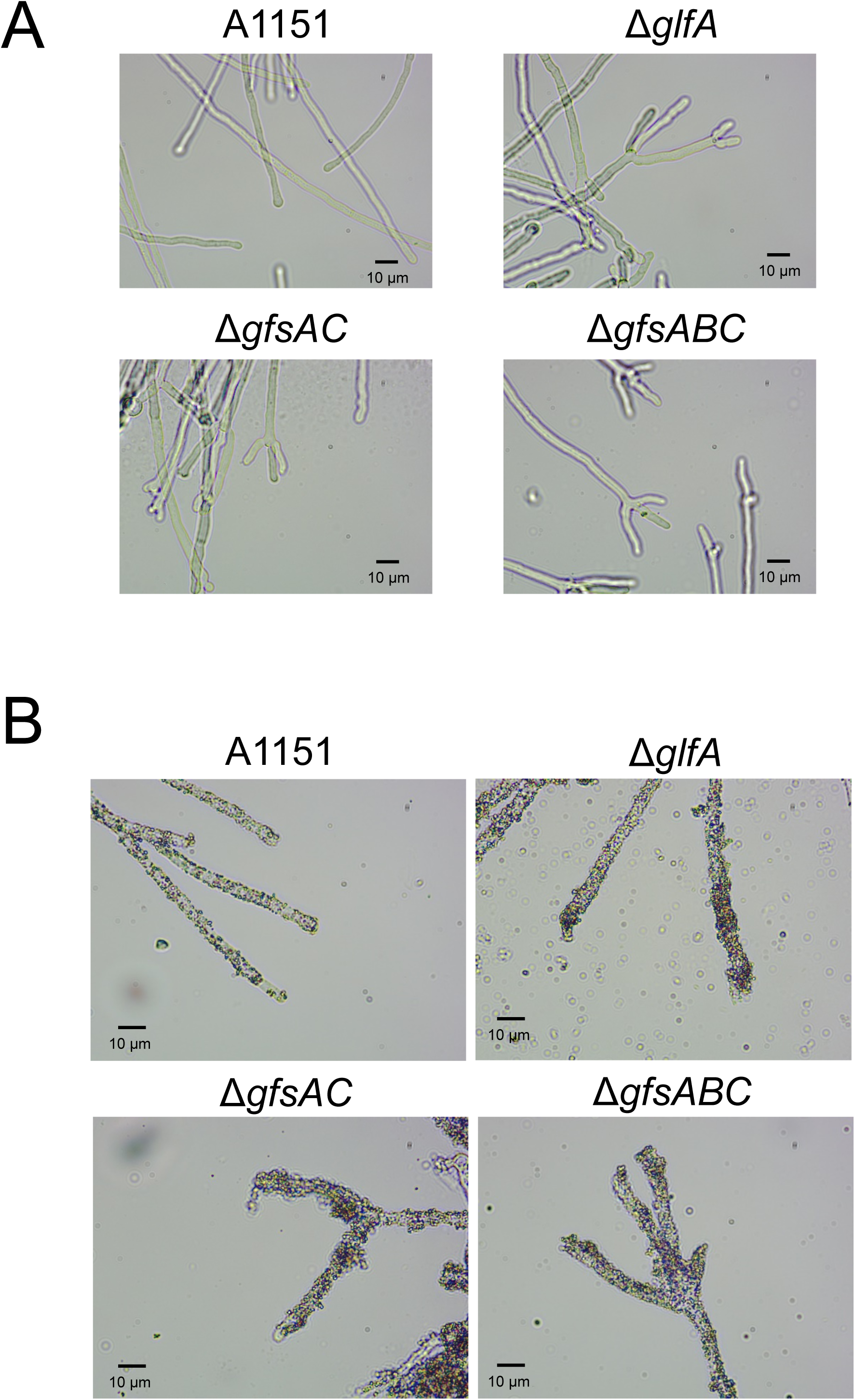
Morphology of the A1151, Δ*glfA*, Δ*gfsAC*, and Δ*gfsABC* strains. (A) Hyphae morphology of the A1151, Δ*glfA*, Δ*gfsAC*, and Δ*gfsABC* strains. (B) Hyphae hydrophobicity of A1151, Δ*glfA*, Δ*gfsAC*, and Δ*gfsABC* strains. Hydrophobicity is indicated by adherence of latex beads to the hyphae.

## DISCUSSION

We previously characterized that GfsA is the β-galactofuranoside β-(1→5)-galactofutanosyltransferase (4). However, the β-(1→5)-galactofuranosyl oligomer synthesized with GfsA could only be confirmed to generate up to 3 sugars in the previous reaction system due to a lack of commercially available UDP-Gal*_f_* (4). In this study, we showed that GfsA could synthesize β-(1→5)-galactofuranosyl oligomers up to lengths of 7 monosaccharides (Fig. 1, *upper panels*). In addition, we showed that GfsB and GfsC also could transfer β-Gal*_f_* to the 5 position of the hydroxy group of the terminal β-galactofuranosyl residue up to 3 and 5 monosaccharides lengths, respectively (Fig. 1, *middle and bottom panels*). AG4, BG2 and CG3 accumulated within the assay of GfsA, GfsB and GfsC, respectively (Fig. 1). This indicates that GfsA is a more suitable enzyme for synthesizing longer β-(1→5)-galactofuranosyl oligomers compared with GfsB and GfsC. This result is consistent with the fact that *gfsA* disruption had greatest impact in the disruptants of *gfs* family genes (Fig. 6). We subsequently proposed structures of the fungal-type- and O-mannose-type galactomannans in the Δ*gfsAC* strain (Fig. 9).

**Figure 8.**
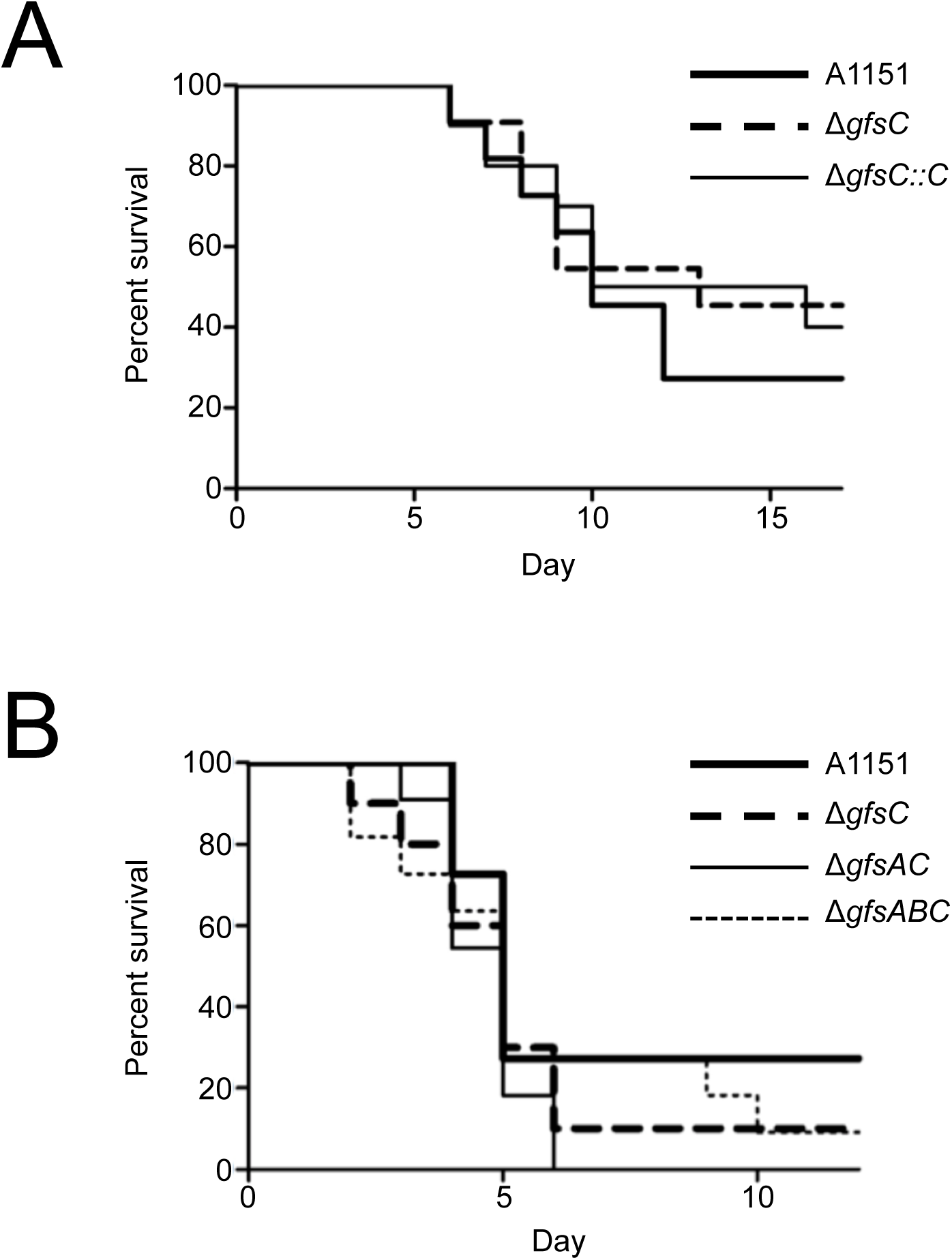
β-(1→5)-galactofuranosyl residues are dispensable for virulence in a mouse model of invasive pulmonary aspergillosis. (A) Infection with A1151, Δ*gfsC* and Δ*gfsC::C* strains. Outbred ICR mice (male; 5 weeks of age; n=11) were immune-compromised via intraperitoneal injection of cyclophosphamide (200 mg/kg) at days −4, −2, 2, and 5. Cortisone acetate was also administrated subcutaneously at a concentration of 200 mg per kg of body weight on day −1. Mice were infected intratracheally with 3.0 × 10^5^ conidia in a volume of 30 μL for each strain (A1151, Δ*gfsC* and Δ*gfsC::C* strains) on day 0. (B) Mouse infection with A1151, Δ*gfsC*, Δ*gfsAC*, and Δ*gfsABC* strains. Outbred ICR mice (male; 5 weeks of age; n=10–11) were immune-compromised via intraperitoneal injection of cyclophosphamide (200 mg/kg mouse) at days −4, −2, 2, and 3. Cortisone acetate was also administrated subcutaneously at a concentration of 200 mg per kg of body weight on day −1. Mice were infected intratracheally with 3.0 × 10^5^ conidia in a volume of 30 μL for each strain (A1151, Δ*gfsC*, Δ*gfsAC*, and Δ*gfsABC*) on day 0.

**Figure 9.**
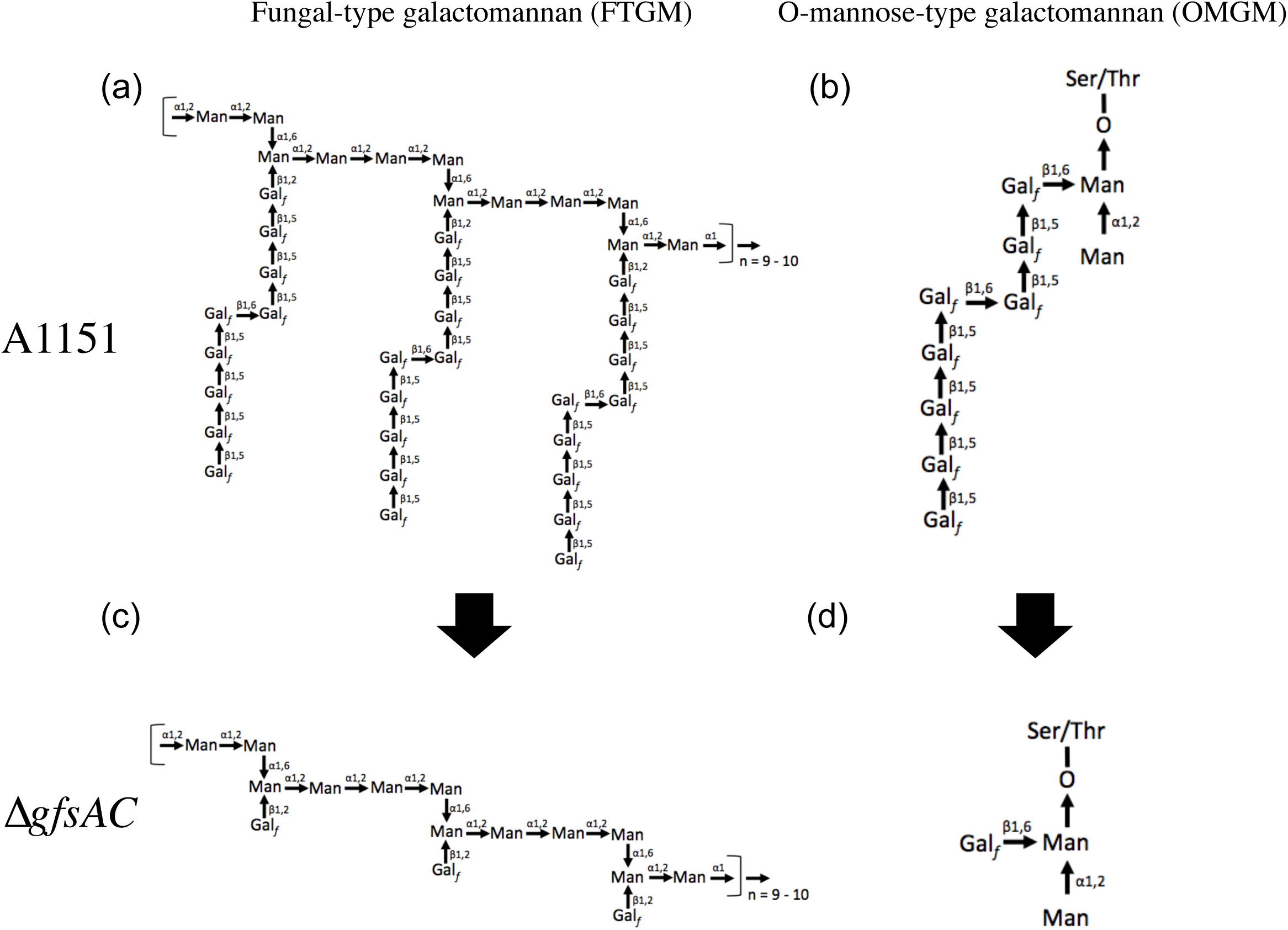
Proposed GM structures in Δ*gfsAC* strain. (A) Typical structure of FTGM, and (B) OMGM in *A. fumigatus*. (C) Proposed structure of FTGM in the Δ*gfsAC* strain. (D) Proposed structure of OMGM in the Δ*gfsAC* strain.

One problematic issue for assaying galactofuranosyltransferases is that the sugar donor UDP-Gal*_f_* is not commercially available. Errey et al. described relatively easily synthesizing UDP-Gal*_f_* using flexible enzymatic and chemo-enzymatic approaches (25). However, obtaining and retaining the chemically unstable UDP-Gal*_f_* remains complicated (26). We thus attempted to establish a galactofuranosyltransferase assay using a continuous reaction of sugar-nucleotide conversion and sugar transfer with UDP-galactopyranose mutase and galactofuranosyltransferase. Rose et al. previously performed a method to detect galactofuranosyltransferase activity via continuous reaction using NADH for the reduction of FAD (27). In our hands galactofuranosylation proceeded even when NADH/NADPH was used instead of SD, but was more efficient with SD versus NADH/NADPH (Fig. S2). This established method could measure galactofuranosyltransferase activity without UDP-Gal*_f_*. In addition, since a sufficient amount of purified product can be separated and purified, this is advantageous for structural analysis of the enzymatic product (Figs. 2 and 3). This method will likely be useful for functional analysis of other galactofuranosyltransferases.

The growth phenotypes of the Δ*gfsA* strain are less severe than that of the Δ*gfsAC*/Δ*gfsABC* strains (Fig. 6, Tables 3 and 4). Deletion of *gfsB* and *gfsC* did not result in any growth defect of *A. fumigatus* (Fig. 6, Tables 3 and 4). Very recently, similar results have been observed within disrupted strains of *gfs* family genes in *A*. *niger* (28), suggesting that existence of a common functional relationships of the *gfs* family proteins for *A. niger* and *A. fumigatus*. We clarified that the phenotypic abnormalities occurring in the Δ*gfsAC* strain were due to defects in β-(1→5)-galactofuranosyl residues via analysis of the sugar chain structure of the Δ*gfsAC* strain.

Several galactofuranosyl sugar chain-absent mutants have been reported in *A*. *fumigatus*; whole galactofurnosyl sugar chains are absent within Δ*glfA* and Δ*glfB* strains (24, 29, 30). These absent Gal*_f_* residues caused decreased growth rates, abnormal hyphae branching, thinner cell walls, increased susceptibility to several antifungal agents and increased adhesive phenotype as compared with the parental strain (24, 29, 30). The phenotypes of the Δ*gfsAC*/Δ*gfsABC* strains were similar in some aspects to the Δ*glfA* strain, but not identical. The latter showed stronger inhibition of hyphal growth and conidia formation compared with Δ*gfsAC*/Δ*gfsABC* (Fig. 6). This is because galactofuranosyl residues are β-(1→5)-linked, β-(1→2)-linked, β-(1→3)-linked, and β-(1→6)-linked sugars (5, 6, 10). These Gal*_f_* residues, except β-(1→5)-galactofuranosyl, are found in glycosyl phosphoinositolceramides (GIPC), FTGM, OMGM, and other sugar chains (5, 6, 10, 31–33) and might be involved in biological events. Only the β-(1→5)-galactofuranosyl residues disappear in the Δ*gfsAC*/Δ*gfsABC* strains, thus it seems reasonable that they exhibit less influence than the Δ*glfA* strain, wherein all Gal*_f_*-containing sugar chain is lost. However, abnormal branching of the hyphae and cell surface hydrophobicity were not significantly different between the Δ*gfsAC*, Δ*gfsABC*, and Δ*glfA* strains (Fig. 7), indicating that GM β-(1→5)-galactofuranosyl residues’ functions are heavily involved in normal polarity of the hyphae and cell surface hydrophobicity.

The presence of the β-(1→6)-galactofuranosyl moiety has been reported in the galactofuran side chain of FTGM and OMGM in *A. fumigatus*. We, therefore, predicted that if the β-(1→5)-galactofuranosyl residues disappeared so would the β-(1→6)-galactofuranosyl residues. However, upon disappearance of the β-(1→5)-galactofuranosyl residues the β-(1→6)-galactofuranosyl residues remained detectable within the ^13^C-NMR of GMs from the Δ*gfsAC* strain (Fig. 4). This strongly suggests the existence of a β-(1→6)-galactofuranosyl oligomer and/or polymer other than the β-(1→6)-galactofuranosyl moiety of the FTGM galactofuran side chain. β-(1→6)-Galactofuranosyl polymer was found in *Fusarium* sp., but not in *A. fumigatus* (34– 36).

In the mouse infection model of invasive aspergillosis, the lack of GM β-(1→5)-galactofuranosyl residues exhibited no significant differences in virulence for the A1151, Δ*gfsAC* and Δ*gfsABC* strains. These findings were consistent with Lamarre’s findings that disruption of *glfA* has no effect on virulence (24). In contrast, Schmalhorst et al. reported that disruption of *glfA* resulted in attenuated virulence in mouse model of invasive aspergillosis (29). Recently, Koch et al. showed that the survival rate of Δ*glfA* strain (Schmalhorst’s strain) decreased slightly more gradually compared with the wild strains using zebrafish embryo model (37). They explained that the attenuated pathogenicity of the Δ*glfA* strain might be caused by decreased germination rate or hyphal growth rate (37). These differences in virulence might be due to varying genetic backgrounds of the strains used or differing protocols of pathogenicity tests, necessitating further detailed analysis to understand the involvement of β-(1→5)-galactofuranosyl sugar chains in pathogenicity.

This study broadens our understanding of the biosynthesis of β-(1→5)-galactofuranosyl residues in *A. fumigatus* and their important role in cell wall formation. However, β-(1→6)-galactofuranosyltransferases that transfer β-galactofuranose to galactofuranosyl residues have not been identified in *A. fumigatus*. Additionally, β-(1→2)-, β-(1→3)-/β-(1→6)-galactofuranosyltransferases transferring β-galactofuranose to mannosyl residues remain unknown. Our findings regarding the biosynthesis of β-(1→5)-galactofuranosyl residue provide important novel insights into the formation of the complex cell wall structure and the virulence of the subphylum Pezisomycotina. Future studies will be needed to identify other galactofuranosyltransferases and clarify the individual functions of each Gal*_f_*-containing oligosaccharide.

## MATERIALS AND METHODS

### Microorganisms and growth conditions

*A. fumigatus* strains used in this study are listed in Table S1. *A. fumigatus* A1160 and A1151 were obtained from the Fungal Genetics Stock Center (FGSC) (38). Strains were grown on minimal medium (MM). Standard transformation procedures for *Aspergillus* strains were used. Plasmids were amplified in *E*. *coli* DH5α. *E. coli* strain Rosetta-gami B (DE3) purchased from Merck Millipore (Merck Millipore, Germany) was used for protein expression. Colony growth rates were measured as described previously (14).

### Construction of GfsB and GfsC expression vector

All PCR reactions were performed using Prime STAR GXL DNA Polymerase (Takara Bio, Otsu, Japan). The pCold™ II (Takara Bio) and pET50b-Amp plasmids for protein expression in *E. coli* were used. The pET50b-Amp is a plasmid constructed by replacing the kanamycin resistance gene of pET50b with an ampicillin resistance gene, and was constructed as follows: DNA region of pET50b except for the kanamycin resistance gene was amplified by PCR using pET50b plasmid as a template for the primer pairs, pET50b-Amp-F and pET50b-Amp-R. The ampicillin resistance gene was amplified by PCR using pET15b plasmid as a template for the primer pairs Amp-gene-F and Amp-gene-R. The obtained DNA fragments were ligated using an In-fusion HD cloning Kit (Takara Bio) to yield pET50b-Amp. Plasmids useful for expression of *gfsB* and *gfsC* were constructed as follows: total RNA was extracted from *A*. *fumigatus* A1160 strain mycelia grown in MM for 18 h using TRIzol Reagent (Thermo Fisher Scientific, MA, USA) according to the manufacturer’s instructions. Single-stranded DNA was synthesized by M-MLV Reverse Transcriptase (NIPPON GENE, Tokyo, Japan) using oligo-dT-18 primers. *gfsB* and *gfsC* were amplified using PCR with single-stranded DNA as a template for primer pairs pET50b-AfGfsB-F and pET50b-AfGfsB-R for *gfsB*, pCold2-AfGfsC-F and pCold2-AfGfsC-R for *gfsC*, respectively. The amplified fragments were inserted into the *Nde* I site of pCold™ II to yield pCold2-AfGfsC, and the *Sma* I site of pET50b-Amp to yield pET50b-Amp-AfGfsB using the In-fusion HD cloning Kit. The constructed plasmids were transformed into Rosetta-gami B (DE3) cells.

### Protein purification, quantification, and electrophoresis

GfsA protein was expressed in Rosetta-gami B (DE3) cells harboring the plasmids pET15b-AfGfsA (4). Protein expression and purification for GfsA and GfsB were performed as described previously (4). Rosetta-gami B (DE3) cells harboring the plasmids pCold2-AfGfsC were used for protein expression of GfsC, which was performed according to the manufacturer’s protocol for the pCold DNA cold-shock expression system. The NusA tag of GfsB was cleaved with a HRV 3C protease at 4°C (Takara Bio) and removed by Ni-agarose. Protein concentrations were determined using the Qubit Protein Assay Kit (Thermo Fisher Scientific), and purified proteins were analyzed by SDS-PAGE to assess purity and molecular weight. Glf protein was obtained with the ASKA clone as previously described (4, 39). Purified Glf was visualized as a band close to the predicted molecular weights of 45.0 kDa (Fig. S1).

### Synthesis of *p*-nitrophenyl β-D-galactofuranoside (pNP-Gal*f*) and 4-methylumbelliferyl β-D-galactofuranoside (4MU-Gal*_f_*)

Para-nitrophenyl β-D-galactofuranoside (pNP-Gal*_f_*) was chemically synthesized as described previously (18, 40) or purchased (Toronto Research Chemicals, Toronto, Canada). 4-methylumbelliferyl β-D-galactofuranoside (4MU-Gal*_f_*) was chemically synthesized as follows (41, 42): 4-methylumbelliferon (50.0 mmol) and BF_3_•Et_2_O (50.0 mmol) was added to a solution of per-benzoylated galactofuranose (10.0 mmol) with 4A molecular sieves in CH_3_CN (50 mL) at 0°C (41). The reaction mixture was stirred at 0°C for 1 h followed by 23°C for 24 h. Next, the mixture was filtered through a Celite pad and the residue was diluted with EtOAc, washed with sat. aq. NaHCO_3_ solution and brine, dried over MgSO_4_, and concentrated *in vacuo* to dryness, producing a mixture of 4-methylumbelliferyl 2,3,4,6-tetra-*O*-benzoyl-β-D-galactofuranoside. A 28% aq. NH_3_ solution was added to the aforementioned mixture in CH_3_OH at 0°C, the resulting solution was stirred at this temperature for 1 h and then at 23°C for 24 h. The reaction solution was concentrated. The target material was purified by silica-gel column chromatography (CHCl_3_:CH_3_OH, 4/1) to give 4-methylumbelliferyl β-D-galactofuranoside (4MU-Gal*_f_*) as a yellow solid (1.80 mmol).

### Continuous enzymatic reaction assay

Standard assays were performed with 1.5 mM 4MU-β-D-Gal*f* acceptor substrate, 40 mM UDP-galactopyranose, purified Glf protein (UDP-galactopyranose mutase from *Escherichia coli*: 15.8 µg), 40 mM sodium dithionite (SD), and purified GfsA (4.5 µg), GfsB (4.5 µg), or GfsC (4.5 µg) proteins in a total reaction volume of 20 µL. The mixtures were incubated at 30°C for 16 h and the reaction was stopped by heat (99°C) for 5 min. The supernatants were analyzed by HPLC with an amino column Shodex Asahipak NH2P-50 4E (250 x 4.6 mm, Showa Denko, Tokyo, Japan) as previously described (4). 4-Methylumbelliferyl and *p*-nitrophenyl derivatives were detected by 300 nm of absorbance. The mass spectra of the enzymatic products of GfsA, GfsB, and GfsC were determined using the Exactive Plus Orbitrap Mass Spectrometer (Thermo Fisher Scientific).

### Construction of Δ*gfsB* and Δ*gfsC* gene disruption strains

*A. fumigatus* A1151/A1160 was used as parental strain (Table S1); *gfsB* were disrupted in the A1151 strain by *ptrA* insertion; *gfsC* was also disrupted in the A1160 strain by *AnpyrG* insertion. DNA fragments for gene disruption were constructed using a “double-joint” PCR method as described previously (43). The 5′- and 3′-flanking regions (approximately 1.0–1.1 kb each) of each gene were PCR amplified from genomic DNA with the following primer pairs (Table S2): AFUB_070620-1/AFUB_070620-2 and AFUB_070620-3/AFUB_070620-4 for *gfsB* disruption; AFUB_067290-1/AFUB_067290-2 and AFUB_067290-3/AFUB_067290-4 for *gfsC*. *ptrA* and *AnpyrG* used as selective markers were amplified using plasmids pPTR-I (Takara Bio) and pSH1 (14) as template, respectively, and the primer pairs ptrA-5/ptrA-6 or pyrG-5/pyrG-6. The three amplified fragments were purified and mixed and a second PCR was performed without specific primers to assemble each fragment, as the overhanging chimeric extensions act as primers. A third PCR was performed with the nested primer pairs AFUB_070620-7/AFUB_070620-8 for *gfsB* or AFUB_067290-7/AFUB_067290-8 for *gfsC* and the products of the second PCR as a template to generate the final deletion construct. The amplified final deletion constructs were purified using the Fast Gene Gel/PCR Extraction Kit (NIPPON GENE) and used directly for transformation. Transformants were grown on MM plates containing 0.6 M KCl as an osmotic stabilizer under appropriate selection conditions and single colonies were isolated twice before further analysis. Disruption of target genes was confirmed by PCR with these primer pairs: AFUB_070620-1/ptrA-R and ptrA-F/AFUB_070620-4 for *gfsB*, AFUB_067290-1/ptrA-R and ptrA-F/AFUB_067290-4 or AFUB_067290-1/pyrG-R and pyrG-F/AFUB_067290-4 for *gfsC* (Fig. S3).

### Construction of the complementary strain of Δ*gfsC* using *gfsC*

*A. fumigatus* Δ*gfsC* strain was used as the parental strain (Table S1). The relevant region of *gfsC* was PCR amplified from genomic DNA using the primer pair AfgfsC-complement-1/AfgfsC-complement-2 (Table S2). The relevant region of the *AnpyrG* was PCR amplified from pSH1 using the primer pair AfgfsC-complement-3/AfgfsC-complement-4 (Table S2). *ptrA* used as selective markers were amplified using the pPTR-I plasmid as a template and the primer pair ptrA-5/ptrA-6. The three amplified fragments were purified and mixed, and a second PCR was performed. A third PCR was performed using the nested primer pair AfgfsA-complement-7/AfgfsA-complement-8 and the products of the second PCR as a template to generate the final DNA construct. Correct replacement of the DNA fragments for gene complementation was confirmed by PCR using the primer pairs AfgfsA-complement-1/ptrA-R and ptrA-F/AfgfsA-complement-4 (Fig. S4).

### Construction of double and triple gene disruption strains

A. fumigatus ΔgfsA was used as a parental strain (Table S1) to construct double and triple gene disruption strains. Genes were disrupted in *A. fumigatus* by *ptrA/hph* insertion; *gfsC* was disrupted in strain Δ*gfsA* by *ptrA* insertion to construct strain Δ*gfsAC*. Next, *gfsB* was disrupted in strain Δ*gfsAC* by *hph* insertion to construct strain Δ*gfsABC*. Primer pairs AFUB_067290-1/AFUB_067290-2(*gfsC*::*ptrA*), AFUB_067290-3(*gfsC*::*ptrA*)/AFUB_067290-4, ptrA-5/ptrA-6 and AFUB_067290-7/AFUB_067290-8 were used to construct a deletion cassette for Δ*gfsC*. Primer pairs AFUB_070620-1/AFUB_070620-2(*gfsB*::*hph*), AFUB_070620-3(*gfsB*::*hph*)/AFUB_070620-4, hph-5/hph-6 and AFUB_070620-7/AFUB_070620-8 were used to construct a deletion cassette for Δ*gfsB*. *hph* was amplified by PCR using pAN7-1 as template (44) and the primers hph-5 and hph-6 (Table S2). Target gene disruption was confirmed using PCR with primer pairs AFUB_096220-1/pyrG-R and pyrG-F/AFUB_096220-4 for *gfsA*, AFUB_070620-1/hph-R and hph-F/AFUB_070620-4 for *gfsB*, and AFUB_067290-1/ptrA-R and ptrA-F/AFUB_067290-4 for *gfsC* (Fig. S5).

### Methylation analysis and Nuclear magnetic resonance (NMR) spectroscopy

GMs were prepared using a previously described method (4, 14). Glycosidic linkage was analyzed using a previously described method (4, 6). NMR experiments were performed as previously described (4, 6, 14). Proton and carbon chemical shifts were referenced relative to internal acetone at δ 2.225 and 31.07 ppm, respectively.

### Analysis of conidiation efficiency and surface adhesion

Conidiation efficiency was analyzed as described previously (14). Hyphal surface adhesion assay was performed as described previously with sight modifications (24, 45). The 0.5-µm diameter polystyrene beads (Sigma) were diluted 1:100 in sterile phosphate buffered saline (PBS). Mycelia were grown for 18 h at 37°C with shaking at 127 rpm in MM liquid medium, harvested into PBS-containing polystyrene beads for 1 h, and then washed five times with PBS. Mycelia images were collected using a microscope equipped with a digital camera.

### Drug susceptibility testing and mouse model of pulmonary aspergillosis

Drug susceptibility was tested in triplicates as described previously (12, 46, 47). The mouse model of pulmonary aspergillosis was generated as per a previously described method with slight modifications (48). In each experiment, A1151, Δ*gfsC*, and Δ*gfsC::C* or A1151, Δ*gfsC*, Δ*gfsAC*, and Δ*gfsABC* strains were used to infect immunosuppressed mice (10 or 11 mice per group). Outbreed male ICR mice were housed in sterile cages (5 or 6 per cage) with sterile bedding and provided with sterile feed and drinking water containing 300 µg/ml tetracycline hydrochloride to prevent bacterial infection. Mice were immunosuppressed with cyclophosphamide (200 mg per kg of body weight), which was intraperitoneally administered on days −4, −2, 2, and 5, or −4, −2, and 3 (day 0: infection). Cortisone acetate (200 mg per kg of body weight) was injected on day −1 for immunosuppression. Mice were infected by intratracheal instillation of 3 × 10^5^ conidia in 30 µl of PBS. Mice were weighed and visually inspected every 24 h from the day of infection. On recording 30% body weight reduction, the mouse was regarded as dead and euthanized. Prism statistical analysis package was used for statistical analysis and survival curve drawing (GraphPad Software Inc., CA, USA).

### Ethics statement

The institutional animal care and use committee of Chiba University approved the animal experiments (Permit Number: DOU28-376 and DOU29-215). All efforts were made to minimize suffering in strict accordance with the principles outlined by the Guideline for Proper Conduct of Animal Experiments.

## ACKNOWLEDGMENTS

This work was supported in part by JSPS KAKENHI grant numbers JP26450106, 18K05418 (to TO) and 17K15492 (to YT), a 2017 Research Grant from the Noda Institute for Scientific Research (to TO) and Joint Usage/Research Program of Medical Mycology Research Center, Chiba University (grant numbers 18-9 and 19-4) (to TO). Strains and plasmids were obtained from the Fungal Genetics Stock Center (Kansas City, MO).

---

**Figure S1.**
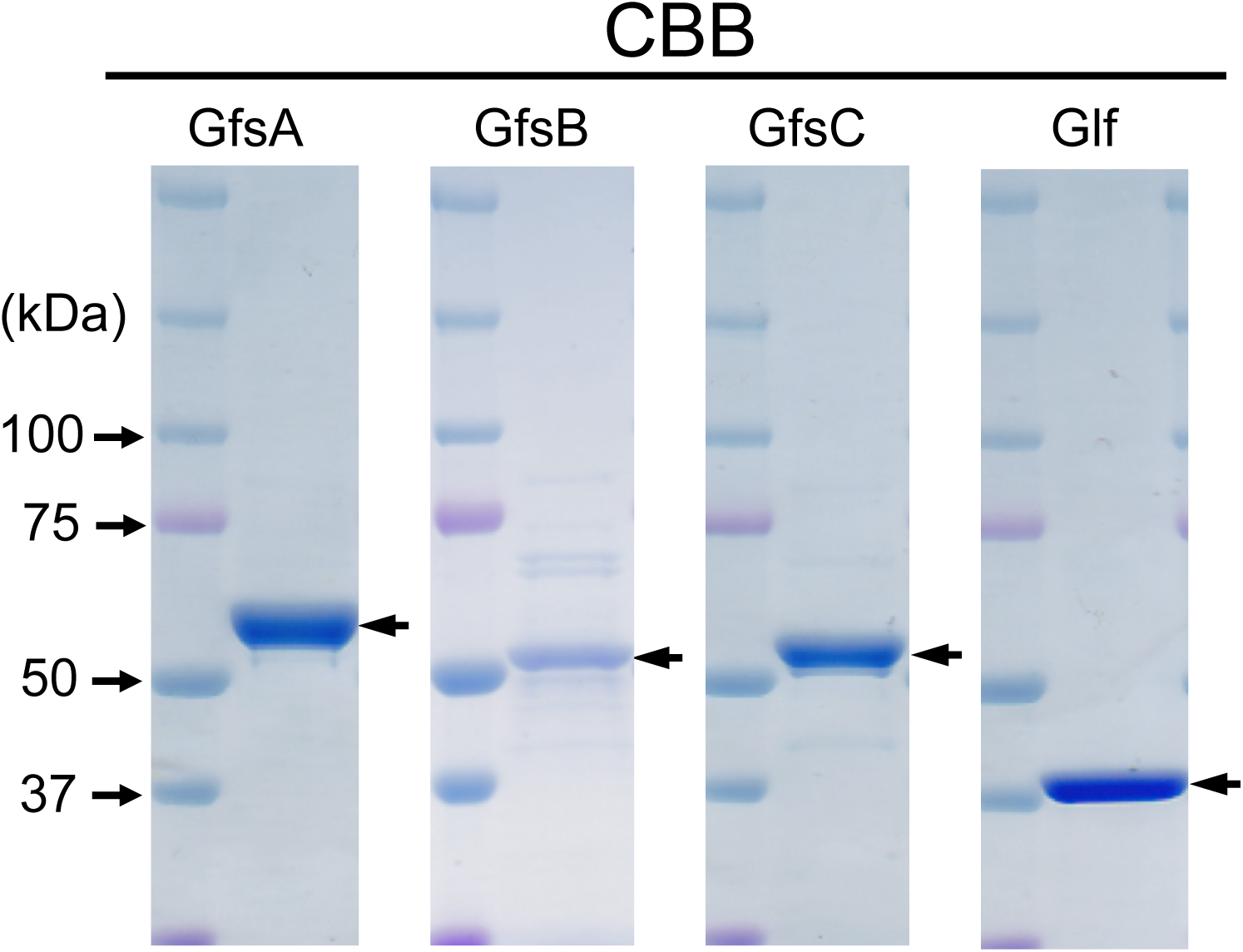
SDS-PAGE analysis of purified recombinant GfsA, GfsB, GfsC, and Glf proteins. Purified recombinant GfsA (5.0 µg), GfsB (3.0 µg), GfsC (5.0 µg), and Glf (5.0 µg) were separated by 5%–20% SDS-PAGE and stained with Coomassie brilliant blue, revealing bands of approximately 57.9 kDa (GfsA), 50.3 kDa (GfsB), 52.0 kDa (GfsC), and 45.0 kDa (Glf).

**Figure S2.**
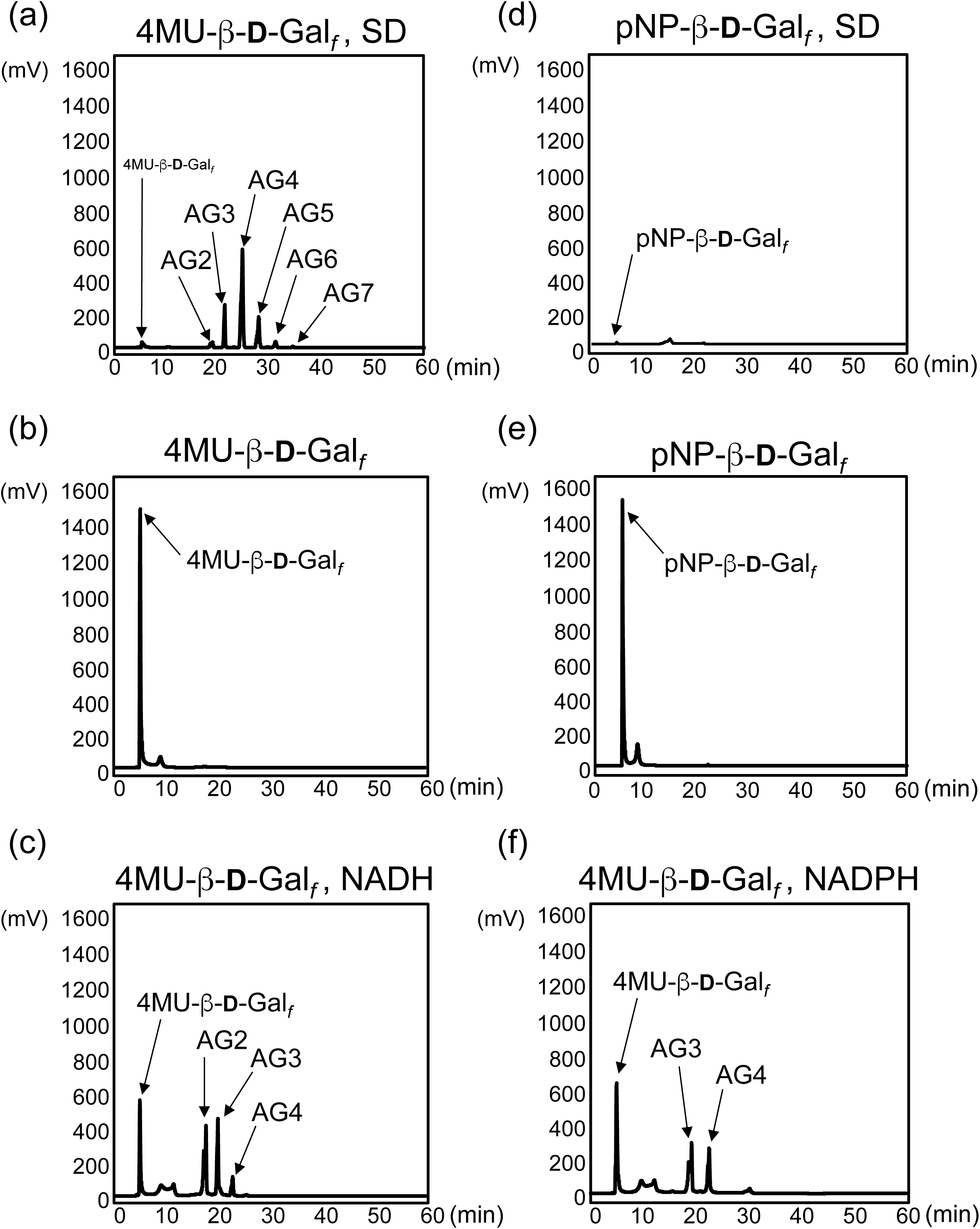
Effects of reducing agent and acceptor substrate on galactofuranosyltransferase assay. Chemically synthesized 4-methylumbelliferyl-β-D-galactofuranoside (4MU-β-D-Gal*_f_*) or *p*-nitrophenyl-β-D-galactofuranoside (pNP-β-D-Gal*_f_*) were used acceptor substrate. The chromatogram of standard galactofuranosyltransferase assay containing the purified GfsA protein (4.5 µg), purified 15.8 µg of Glf protein, 40 mM UDP-Gal*_p_*, 1 mM Mn^2+^, 1.5 mM 4MU-β-D-Gal*_f_*, and 40 mM sodium dithionite (SD) at 30°C for 16 h (a); 40 mM SD was omitted from the standard assay (b); 40 mM NADH was added to the standard assay instead of 40 mM SD (c), pNP-β-D-Gal*_f_* was used as an acceptor substrate instead of 4MU-β-D-Gal*_f_* (d), or pNP-β-D-Gal*_f_* instead of 4MU-β-D-Gal*_f_* and 40 mM SD was omitted from the standard assay (e); and 40 mM NADPH was added to the standard assay instead of 40 mM SD (f).

**Figure S3.**
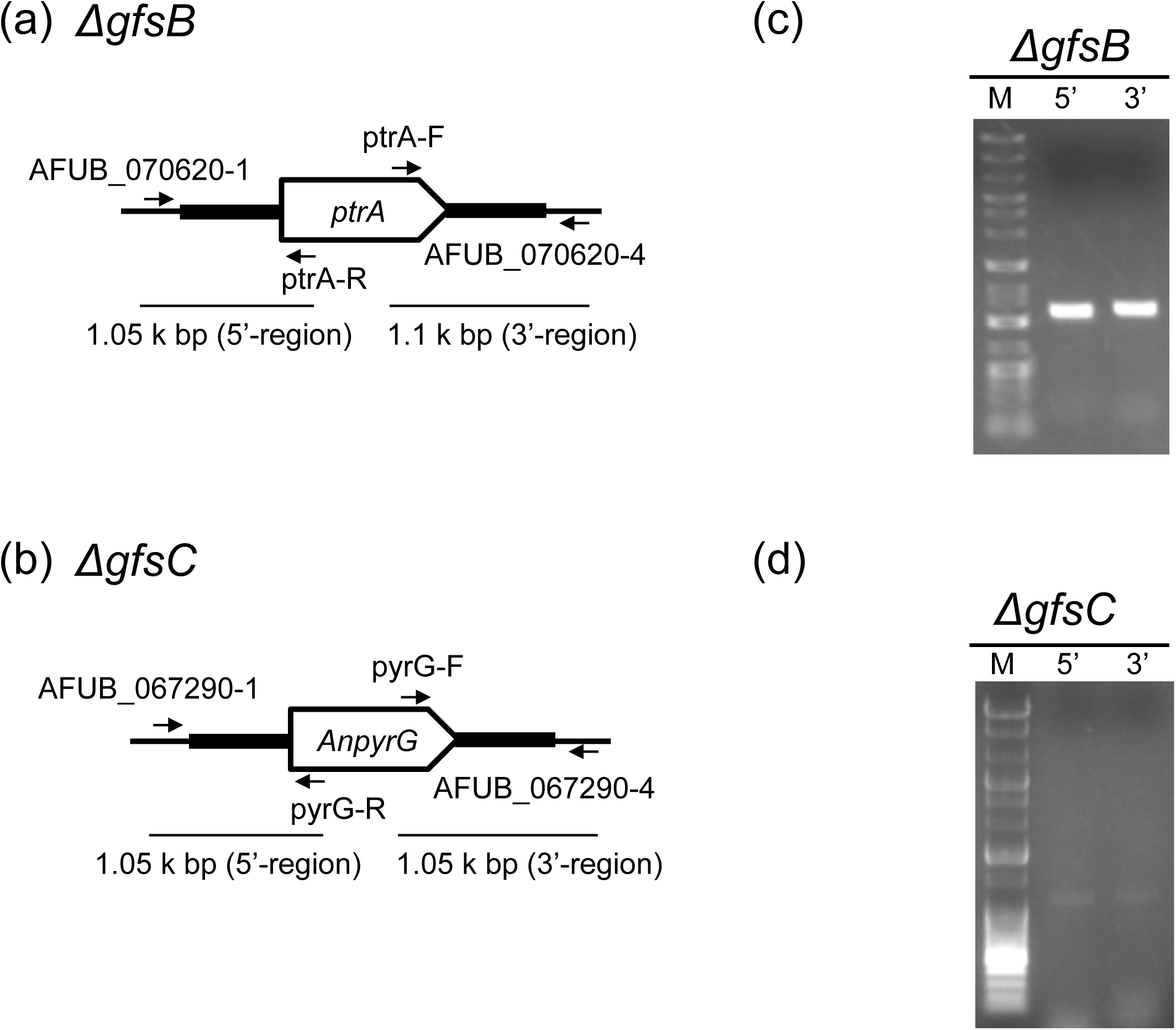
Construction of the strains Δ*gfsB*, Δ*gfsC* (*ptrA*), and Δ*gfsC* (*AnpyrG*). Chromosomal maps of strains Δ*gfsB* (a) and Δ*gfsC* (b), and primers used for confirmation. The positions of the primers are indicated by arrows. Electrophoretic analyses of products amplified by PCR using the primer pairs AFUB_070620-1/ptrA-R (5′-region) and ptrA-F/AFUB_070620-4 (3′-region) for Δ*gfsB* (c), and AFUB_067290-1/pyrG-R (5′-region) and pyrG-F/AFUB_067290-4 (3′-region) for Δ*gfsC* (d). M: DNA size markers; Gene Ladder Wide 2 (Nippon Gene, Tokyo, Japan).

**Figure S4.**
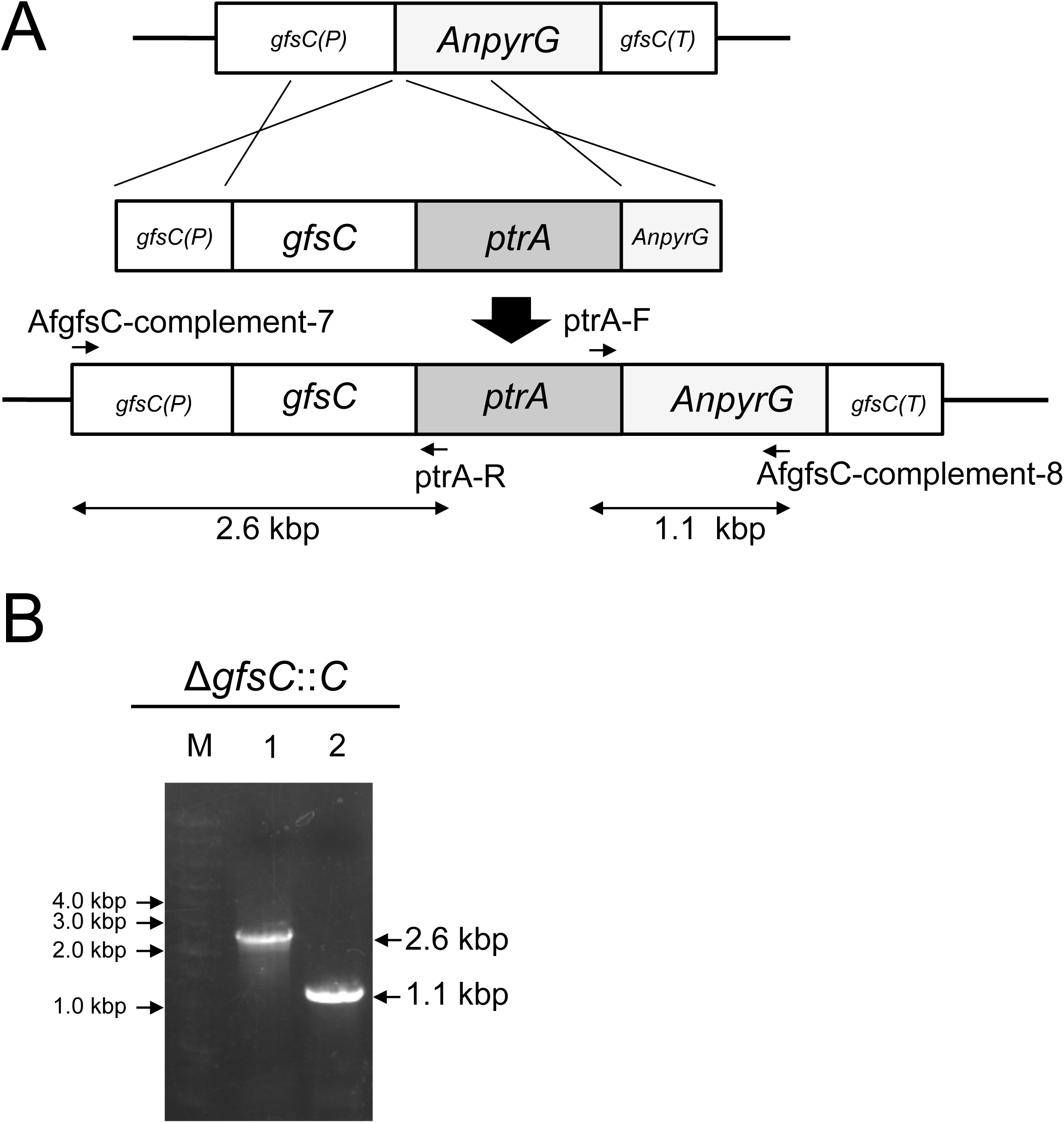
Construction of the Δ*gfsC* complementary strains Δ*gfsC::C*. (a) Schematic representation of Δ*gfsC* complementation with *gfsC*. *gfsC* (P), *gfsC* promoter; *gfsC* (T), *gfsC* terminator; *gfsC*, open reading frame of *gfsC*. The positions of the primers are indicated by arrows. (b) Confirmation of correct recombination of *gfsC* using PCR analysis. Electrophoretic analysis of products amplified by PCR are shown. M, DNA size markers; Gene Ladder Wide 2; Lane 1, DNA fragment (2.6 kbp) amplified using PCR and the primers AfgfsC-complement-7 and ptrA-R; lane 2, DNA fragment (1.1 kb) amplified using PCR and the primers ptrA-F and AfgfsC-complement-8.

**Figure S5.**
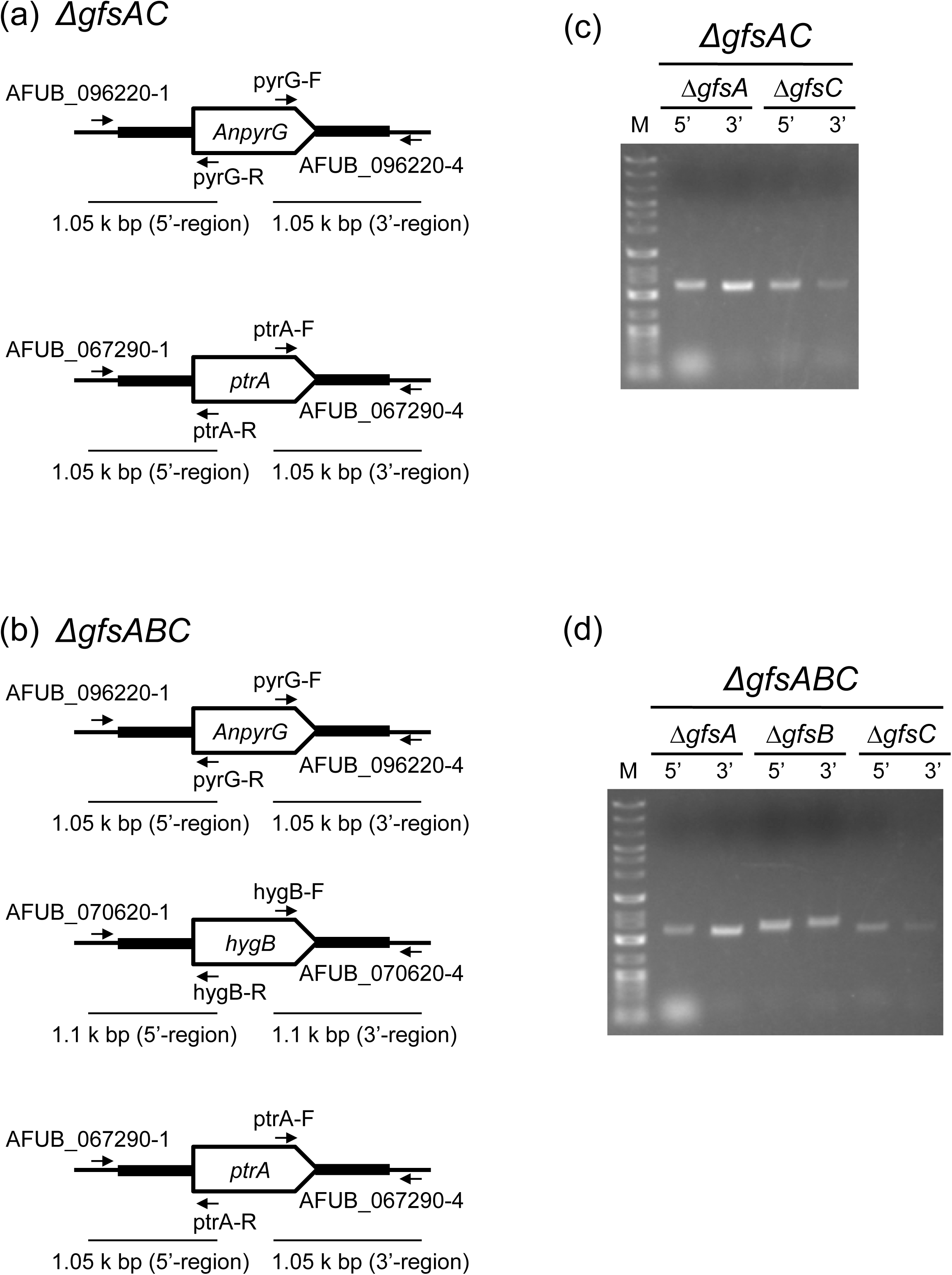
Construction of the strains Δ*gfsAC* and Δ*gfsABC*. (a) Chromosomal maps of strains Δ*gfsAC* (a) and Δ*gfsABC* (b), and primers used for confirmation. The positions of the primers are indicated by arrows. Electrophoretic analyses of products amplified by PCR using the primer pairs AFUB_096220-1/pyrG-R (5′ region of *gfsA*), pyrG-F/AFUB_096220-4 (3′ region of *gfsA*), AFUB_067290-1/ptrA-R (5′ region of *gfsC*), and ptrA-F/AFUB_067290-4 (3′ region of *gfsC*), for Δ*gfsAC* (c), AFUB_096220-1/pyrG-R (5′ region of *gfsA*, pyrG-F/AFUB_096220-4 (3′ region of *gfsA*), AFUB_070620-1/hygB-R (5′ region of *gfsB*), hygB-F/AFUB_070620-4 (3′ region of *gfsB*), AFUB_067290-1/ptrA-R (5′ region of *gfsC*), and ptrA-F/AFUB_067290-4 (3′ region of *gfsC*), for Δ*gfsABC* (d). M: DNA size markers; Gene Ladder Wide 2.

**Table S1.** Aspergillus strains used in this study.

| Strains | Genotype | Source of reference |
| --- | --- | --- |
| *A. fumigatus* A1151 | *KU80::pyrG* | da Silva Ferreira, 2006; FGSC |
| *A. fumigatus* A1160 | *KU80::pyrG, pyrG<sup>-</sup>* | da Silva Ferreira, 2006; FGSC |
| *A. fumigatus* Δ*glfA* | *KU80::AfpyrG, glfA::ptrA* | Komachi, 2013 (ref) |
| *A. fumigatus* Δ*gfsA* | *KU80::AfpyrG, pyrG<sup>-</sup>, AfgfsA::AnpyrG* | Komachi, 2013 (ref) |
| *A. fumigatus* Δ*gfsB* | *KU80::AfpyrG, AfgfsB::ptrA* | This study |
| *A. fumigatus* Δ*gfsC* | *KU80::AfpyrG, pyrG<sup>-</sup>, AfgfsC::AnpyrG* | This study |
| *A. fumigatus* Δ*gfsAC* | *KU80::AfpyrG, pyrG<sup>-</sup>, AfgfsA::AnpyrG, AfgfsC::ptrA* | This study |
| *A. fumigatus* Δ*gfsABC* | *KU80::AfpyrG, pyrG<sup>-</sup>, AfgfsA::AnpyrG, AfgfsB::hph, AfgfsC::ptrA* | This study |
| *A. fumigatus* Δ*gfsA::A* | *KU80::pyrG, AfgfsA::AnpyrG, ΔAfgfsA::AfgfsA-*ptrA** | Katafuchi, 2017 (ref) |
| *A. fumigatus* Δ*gfsC::C* | *KU80::AfpyrG, pyrG<sup>-</sup>, AfgfsC::AnpyrG, ΔAfgfsC::AfgfsC-*ptrA** | This study |

**Table S2.**
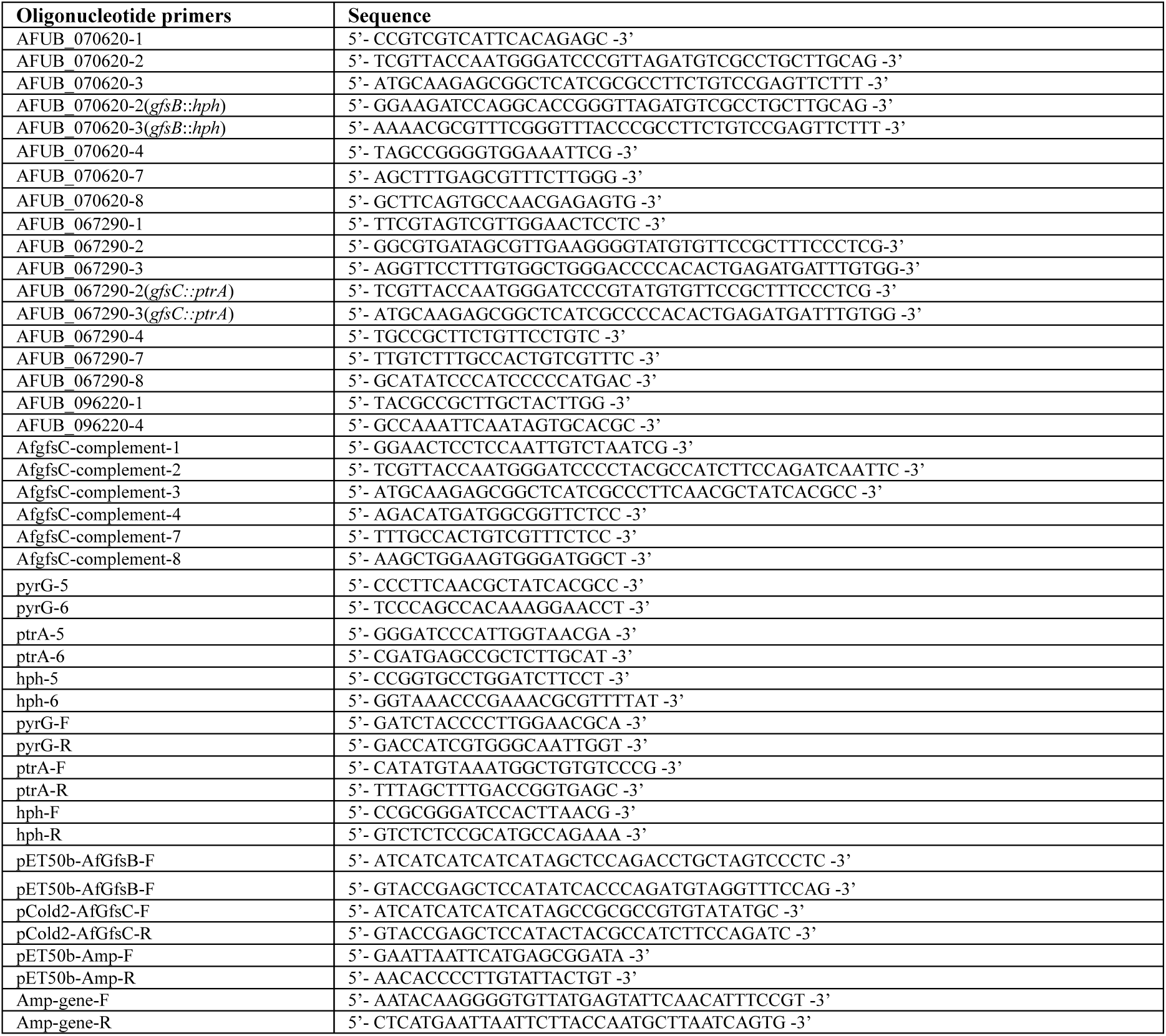
Oligonucleotides used in this study.

## REFERENCES

1. Latgé JP, Beauvais A, Chamilos G. 2017. The cell wall of the human fungal pathogen *Aspergillus fumigatus*: biosynthesis, organization, immune response, and virulence. Annu Rev Microbiol 71:99–116. https://doi.org/10.1146/annurev-micro-030117-020406.

2. Gow NAR, Latge JP, Munro CA. 2017. The Fungal cell wall: structure, biosynthesis, and function. Microbiol Spectr 5 https://doi.org/10.1128/microbiolspec.FUNK-0035-2016.

3. Oka T. 2018. Biosynthesis of galactomannans found in filamentous fungi belonging to Pezizomycotina. Biosci Biotechnol Biochem 82:183–191. https://doi.org/10.1080/09168451.2017.1422383.

4. Katafuchi Y, Li Q, Tanaka Y, Shinozuka S, Kawamitsu Y, Izumi M, Ekino K, Mizuki K, Takegawa K, Shibata N, Goto M, Nomura Y, Ohta K, Oka T. 2017. GfsA is a β1,5-galactofuranosyltransferase involved in the biosynthesis of the galactofuran side chain of fungal-type galactomannan in *Aspergillus fumigatus*. Glycobiology 27:568–581. https://doi.org/10.1093/glycob/cwx028.

5. Latgé JP, Kobayashi H, Debeaupuis JP, Diaquin M, Sarfati J, Wieruszeski JM, Parra E, Bouchara JP, Fournet B. 1994. Chemical and immunological characterization of the extracellular galactomannan of *Aspergillus fumigatus*. Infect Immun 62:5424–5433.

6. Kudoh A, Okawa Y, Shibata N. 2015. Significant structural change in both *O*-and *N*-linked carbohydrate moieties of the antigenic galactomannan from *Aspergillus fumigatus* grown under different culture conditions. Glycobiology 25:74–87. https://doi.org/10.1093/glycob/cwu091.

7. Costachel C, Coddeville B, Latgé JP, Fontaine T. 2005. Glycosylphosphatidylinositol-anchored fungal polysaccharide in *Aspergillus fumigatus*. J Biol Chem 280:39835–39842. https://doi.org/10.1074/jbc.m510163200.

8. Muszkieta L, Fontaine T, Beau R, Mouyna I, Vogt MS, Trow J, Cormack BP, Essen LO, Jouvion G, Latgé JP. 2019. The Glycosylphosphatidylinositol-anchored *DFG* family is essential for the insertion of galactomannan into the β-(1,3)-glucan-chitin core of the cell wall of *Aspergillus fumigatus*. mSphere 4 pii: e00397–19. https://doi.org/10.1128/mSphere.00397-19.

9. Oka T, Hamaguchi T, Sameshima Y, Goto M, Furukawa K. 2004. Molecular characterization of protein O-mannosyltransferase and its involvement in cell-wall synthesis in Aspergillus nidulans. Microbiology 150:1973–1982. https://doi.org/10.1099/mic.0.27005-0.

10. Tefsen B, Ram AF, van Die I, Routier FH. 2012. Galactofuranose in eukaryotes: aspects of biosynthesis and functional impact. Glycobiology 22:456–469. https://doi.org/10.1093/glycob/cwr144.

11. Oka T, Goto M 2016. Biosynthesis of galactofuranose-containing glycans in filamentous fungi. Trends Glycosci Glycotechnol 28:E39–E45. https://doi.org/10.4052/tigg.1428.1E.

12. Onoue T, Tanaka Y, Hagiwara D, Ekino K, Watanabe A, Ohta K, Kamei K, Shibata N, Goto M, Oka T. 2018. Identification of two mannosyltransferases contributing to biosynthesis of the fungal-type galactomannan α-core-mannan structure in *Aspergillus fumigatus*. Sci Rep 8:16918. https://doi.org/10.1038/s41598-018-35059-2.

13. Henry C, Li J, Danion F, Alcazar-Fuoli L, Mellado E, Beau R, Jouvion G, Latgé JP, Fontaine T. 2019. Two KTR mannosyltransferases are responsible for the biosynthesis of cell wall mannans and control polarized growth in *Aspergillus fumigatus*. mBio 10: e02647–18. https://doi.org/10.1128/mBio.02647-18.

14. Komachi Y, Hatakeyama S, Motomatsu H, Futagami T, Kizjakina K, Sobrado P, Ekino K, Takegawa K, Goto M, Nomura Y, Oka T. 2013. GfsA encodes a novel galactofuranosyltransferase involved in biosynthesis of galactofuranose antigen of *O*-glycan in *Aspergillus nidulans* and *Aspergillus fumigatus*. Mol Microbiol 90:1054–1073. https://doi.org/10.1111/mmi.12416.

15. Nassau PM, Martin SL, Brown RE, Weston A, Monsey D, McNeil MR, Duncan K. 1996. Galactofuranose biosynthesis in *Escherichia coli* K-12: identification and cloning of UDP-galactopyranose mutase. J Bacteriol 178:1047–1052. https://doi.org/10.1128/jb.178.4.1047-1052.1996.

16. Lee R, Monsey D, Weston A, Duncan K, Rithner C, McNeil M. 1996. Enzymatic synthesis of UDP-galactofuranose and an assay for UDP-galactopyranose mutase based on high-performance liquid chromatography. Anal Biochem 242:1–7. https://doi.org/10.1006/abio.1996.0419.

17. Varela O, Marino C, de Lederkremer RM. 1986. Synthesis of *p*-nitrophenyl beta-D-galactofuranoside. A convenient substrate for beta-galactofuranosidase. Carbohydr Res 155:247–251. https://doi.org/10.1016/S0008-6215(00)90153-8.

18. Ota R, Okamoto Y, Vavricka CJ, Oka T, Matsunaga E, Takegawa K, Kiyota H, Izumi M. 2019. Chemo-enzymatic synthesis of *p*-nitrophenyl β-D-galactofuranosyl disaccharides from *Aspergillus* sp. fungal-type galactomannan. Carbohydr Res 473:99–103. https://doi.org/10.1016/j.carres.2019.01.005.

19. Wasmuth CR, Edwards C, Hutcherson R. 1964. Participation of the SO_2_^-^ Radical Ion in the Reduction of *p*-Nitrophenol by Sodium Dithionite. J Phys Chem 68: 423–425. https://doi.org/10.1021/j100784a510.

20. Shibata N, Saitoh T, Tadokoro Y, Okawa Y. 2009. The cell wall galactomannan antigen from *Malassezia furfur* and *Malassezia pachydermatis* contains beta-1,6-linked linear galactofuranosyl residues and its detection has diagnostic potential. Microbiology 155:3420–3429. https://doi.org/10.1099/mic.0.029967-0.

21. Rose NL, Zheng RB, Pearcey J, Zhou R, Completo GC, Lowary TL. 2008. Development of a coupled spectrophotometric assay for GlfT2, a bifunctional mycobacterial galactofuranosyltransferase. Carbohydr Res 343:2130–2139. https://doi.org/10.1016/j.carres.2008.03.023.

22. May JF, Splain RA, Brotschi C, Kiessling LL. A tethering mechanism for length control in a processive carbohydrate polymerization. 2009 Proc Natl Acad Sci U S A 106:11851–11856. https://doi.org/10.1073/pnas.0901407106.

23. Shibata N, Okawa Y. 2011. Chemical structure of beta-galactofuranose containing polysaccharide and *O*-linked oligosaccharides obtained from the cell wall of pathogenic dematiaceous fungus *Fonsecaea pedrosoi*. Glycobiology 21:69–81. https://doi.org/10.1093/glycob/cwq132.

24. Lamarre C, Beau R, Balloy V, Fontaine T, Wong Sak Hoi J, Guadagnini S, Berkova N, Chignard M, Beauvais A, Latgé JP. 2009 Galactofuranose attenuates cellular adhesion of Aspergillus fumigatus. Cell Microbiol 11:1612–1623. https://doi.org/10.1111/j.1462-5822.2009.01352.x.

25. Errey JC, Mukhopadhyay B, Kartha KP, Field RA. 2004. Flexible enzymatic and chemo-enzymatic approaches to a broad range of uridine-diphospho-sugars. Chem Commun (Camb) 7:2706–2707. https://doi.org/10.1039/b410184g.

26. Tsvetkova YE, Nikolaev AV. 2000. The first chemical synthesis of UDP-α-D-galactofuranose. J Chem Soc Perkin Trans 1:889–891. https://doi.org/10.1039/B000210K.

27. Rose NL, Completo GC, Lin SJ, McNeil M, Palcic MM, Lowary TL. 2006. Expression, purification, and characterization of a galactofuranosyltransferase involved in *Mycobacterium tuberculosis* arabinogalactan biosynthesis. J Am Chem Soc 128:6721–6729. https://doi.org/10.1021/ja058254d.

28. Arentshorst M, de Lange D, Park J, Lagendijk EL, Alazi E, van den Hondel CAMJJ, Ram AFJ. 2019. Functional analysis of three putative galactofuranosyltransferases with redundant functions in galactofuranosylation in Aspergillus niger. Arch Microbiol https://doi.org/10.1007/s00203-019-01709-w.

29. Schmalhorst PS, Krappmann S, Vervecken W, Rohde M, Müller M, Braus GH, Contreras R, Braun A, Bakker H, Routier FH. 2008. Contribution of galactofuranose to the virulence of the opportunistic pathogen *Aspergillus fumigatus*. Eukaryot Cell 2008 7:1268–1277. https://doi.org/10.1128/EC.00109-08.

30. Engel J, Schmalhorst PS, Dörk-Bousset T, Ferrières V, Routier FH. 2009. A single UDP-galactofuranose transporter is required for galactofuranosylation in *Aspergillus fumigatus*. J Biol Chem 284:33859–33868. https://doi.org/10.1074/jbc.M109.070219.

31. Toledo MS, Levery SB, Bennion B, Guimaraes LL, Castle SA, Lindsey R, Momany M, Park C, Straus AH, Takahashi HK. 2007. Analysis of glycosylinositol phosphorylceramides expressed by the opportunistic mycopathogen *Aspergillus fumigatus*. J Lipid Res 48:1801–1824. https://doi.org/10.1194/jlr.m700149-jlr200.

32. Guimarães LL, Toledo MS, Ferreira FA, Straus AH, Takahashi HK. 2014. Structural diversity and biological significance of glycosphingolipids in pathogenic and opportunistic fungi. Front Cell Infect Microbiol 4:138. https://doi.org/10.3389/fcimb.2014.00138.

33. Kotz A, Wagener J, Engel J, Routier F, Echtenacher B, Pich A, Rohde M, Hoffmann P, Heesemann J, Ebel F. 2010. The *mitA* gene of *Aspergillus fumigatus* is required for mannosylation of inositol-phosphorylceramide, but is dispensable for pathogenicity. Fungal Genet Biol 47:169–178. https://doi.org/10.1016/j.fgb.2009.10.001.

34. Iwahara S, Maeyama T, Mishima T, Jikibara T, Takegawa K, Iwamoto H 1992. Studies on the uronic acid-containing glycoproteins of Fusarium sp. M7-1: IV. Isolation and identification of four novel oligosaccharide units derived from the acidic polysaccharide chain. J Biochem 112:355–359. https://doi.org/10.1093/oxfordjournals.jbchem.a123905.

35. Iwahara S, Mishima T, Ramli N, Takegawa K. 1996. Degradation of beta 1 -->6 galactofuranoside linkages in the polysaccharide of Fusarium sp. M7-1 by endo-beta-galactofuranosidase from Bacillus sp. Biosci Biotechnol Biochem 60:957–961. https://doi.org/10.1271/bbb.60.957.

36. Chen YL, Mao WJ, Tao HW, Zhu WM, Yan MX, Liu X, Guo TT, Guo T. 2015. Preparation and characterization of a novel extracellular polysaccharide with antioxidant activity, from the mangrove-associated fungus *Fusarium oxysporum*. Mar Biotechnol (NY) 17:219–228. https://doi.org/10.1007/s10126-015-9611-6.

37. Koch BEV, Hajdamowicz NH, Lagendijk E, Ram AFJ, Meijer AH. 2019. *Aspergillus fumigatus* establishes infection in zebrafish by germination of phagocytized conidia, while *Aspergillus niger* relies on extracellular germination. Sci Rep. 9:12791. https://doi.org/10.1038/s41598-019-49284-w.

38. da Silva Ferreira ME, Kress MR, Savoldi M, Goldman MH, Härtl A, Heinekamp T, Brakhage AA, Goldman GH. 2006. The *akuB*(KU80) mutant deficient for nonhomologous end joining is a powerful tool for analyzing pathogenicity in *Aspergillus fumigatus*. Eukaryot Cell 5:207–211. https://doi.org/10.1128/ec.5.1.207-211.2006.

39. Kitagawa M, Ara T, Arifuzzaman M, Ioka-Nakamichi T, Inamoto E, Toyonaga H, Mori H. 2005. Complete set of ORF clones of *Escherichia coli* ASKA library (a complete set of *E. coli* K-12 ORF archive): unique resources for biological research. DNA Res 12:291–299. https://doi.org/10.1093/dnares/dsi012.

40. Matsunaga E, Higuchi Y, Mori K, Yairo N, Oka T, Shinozuka S, Tashiro K, Izumi M, Kuhara S, Takegawa K. 2015. Identification and characterization of a novel galactofuranose-specific β-D-galactofuranosidase from *Streptomyces* species. PLoS One 10:e0137230. http://dx.doi.org/10.1371/journal.pone.0137230

41. D’Accorso NB, Thiel IME, Schüller M. 1983. Proton and C-13 nuclear magnetic resonance spectra of some benzoylated aldohexoses. Carbohydr. Res 124:177–184. https://doi.org/10.1016/0008-6215(83)88453-5.

42. de Lederkremer RM, Nahmad VB, Varela O. 1994. Synthesis of α-ᴅ-Galactofuranosyl Phosphate. J Org Chem 59, 690–692. http://dx.doi.org/10.1021/jo00082a037

43. Yu JH, Hamari Z, Han KH, Seo JA, Reyes-Domínguez Y, Scazzocchio C. 2004. Double-joint PCR: a PCR-based molecular tool for gene manipulations in filamentous fungi. Fungal Genet Biol. 41:973–981. https://doi.org/10.1016/j.fgb.2004.08.001.

44. Punt PJ, Oliver RP, Dingemanse MA, Pouwels PH, van den Hondel CA. 1987. Transformation of *Aspergillus* based on the hygromycin B resistance marker from *Escherichia coli*. Gene 56:117–124. https://doi.org/10.1016/0378-1119(87)90164-8.

45. Alam MK, van Straaten KE, Sanders DA, Kaminskyj SG. 2014. *Aspergillus nidulans* cell wall composition and function change in response to hosting several *Aspergillus fumigatus* UDP-galactopyranose mutase activity mutants. PLoS One 9:e85735. https://doi.org/10.1371/journal.pone.0085735.

46. Wayne, P. A. 2008. Clinical and laboratory standards institute reference method for broth dilution antifungal susceptibility testing of filamentous fungi. CLSI. Approved standard-second edition clinical and laboratory standards institute document M38–A2.

47. Kikuchi K, Watanabe A, Ito J, Oku Y, Wuren T, Taguchi H, Yarita K, Muraosa Y, Yahiro M, Yaguchi T, Kamei K. 2014. Antifungal susceptibility of *Aspergillus fumigatus* clinical isolates collected from various areas in Japan. J Infect Chemother 20:336–338. https://doi.org/10.1016/j.jiac.2014.01.003.

48. Hagiwara D, Miura D, Shimizu K, Paul S, Ohba A, Gonoi T, Watanabe A, Kamei K, Shintani T, Moye-Rowley WS, Kawamoto S, Gomi K. 2017. A Novel Zn_2_-Cys_6_ Transcription Factor AtrR Plays a key role in an azole resistance mechanism of *Aspergillus fumigatus* by co-regulating *cyp51A* and *cdr1B* expressions. PLoS Pathog 13:e1006096. https://doi.org/10.1371/journal.ppat.1006096.

